# Distinct Sox9 single-molecule dynamics characterize adult differentiation and fetal-like reprogrammed states in intestinal organoids

**DOI:** 10.1101/2025.05.16.654343

**Authors:** Nike Walther, Sathvik Anantakrishnan, Gina M. Dailey, Anna C. Maurer, Claudia Cattoglio

**Affiliations:** Department of Molecular and Cell Biology, Li Ka Shing Center for Biomedical and Health Sciences, California Institute for Regenerative Medicine (CIRM) Center of Excellence, University of California, Berkeley, Berkeley, California 94720, USA; Current affiliation: Department of Genetics, University of Bayreuth, 95440 Bayreuth, Germany; Biophysics Graduate Group, University of California, Berkeley, Berkeley, California 94720, USA; Current affiliation: Department of Biophysics, University of Michigan, Ann Arbor, Michigan 48109, USA; Current affiliation: Department of Microbiology and Immunology, University of Michigan Medical School, Ann Arbor, Michigan 48109, USA; Howard Hughes Medical Institute, Berkeley, California 94720, USA

**Keywords:** Transcription factor, Sox9, Mouse small intestinal organoid, 2D enteroid monolayer culture, Automated live-cell single-molecule tracking, Cellular feature extraction and protein diffusion correlation, Cellular protein diffusion and self-association, Differentiation, Fetal-like reversion, Cell state transition, Transcription factor dosage

## Abstract

Transcription factors (TFs) mediate gene expression changes during differentiation and development. However, how TF biophysical properties and abundance dynamically regulate specific cell state transitions remains poorly understood.

Using automated live-cell single-molecule tracking (SMT) in intestinal organoid models, we revealed an expression level-independent decrease in the fraction of immobile Sox9 molecules during differentiation from ∼48% to ∼38%, largely dependent on DNA binding. Strikingly, long-term Sox9 overexpression caused organoids to transition from budding to spheroid morphology accompanied by increased proliferation and a loss in gene expression signatures for intestinal identity and function. In this fetal-like reprogrammed state, a larger fraction of partially self-interacting Sox9 molecules (∼61%) binds to DNA.

Our results suggest context-dependent Sox9 single-molecule dynamics during adult intestinal differentiation and fetal-like reversion in consequence to long-term Sox9 overexpression. Our work underpins the power of our automated live-cell SMT framework to generate testable hypotheses towards unraveling molecular mechanisms underlying tissue-level phenotypes.

**Highlights:**

- Heterogenous diffusion behavior of Sox9 across intestinal differentiation states
- Concentration-independent decrease in chromatin-bound Sox9 upon differentiation
- Sox9 overexpression leads to fetal-like reversion and more immobile Sox9 molecules
- Automated SMT in 2D monolayers reveals TF dynamics underlying organoid phenotypes

## Introduction

Gene expression programs are rewired during tissue development and homeostasis to ensure the formation and maintenance of healthy organs. This is accomplished by the differentiation of stem cells into various specialized cell types in a highly spatiotemporally controlled manner. One such regulatory layer is provided by lineage-specific transcription factors (TFs)^1^. Acting downstream of signaling pathways, TFs bind with co-factors to *cis*-regulatory elements of target genes, inducing or repressing their expression^2^. Misexpression of TFs conferring cell fate decisions can lead to cellular reprogramming and is associated with disease, including cancer^3,4^. It is thus crucial that lineage TFs are expressed in the correct tissue place at the correct dose. However, it remains poorly understood how the abundance and biophysical properties of cell fate-determining TFs change during stem cell differentiation to mature cell types. It is further unclear what TF dosage is tolerated for faithful differentiation and tissue formation, and whether TF overexpression alters their molecular dynamics. To address these questions, TF abundance and single-molecule dynamics need to be probed in differentiating, multicellular systems, necessitating the choice of a model system that recapitulates *in vivo* differentiation trajectories and is amenable to live-cell single-molecule imaging across scales.

The mammalian intestine as the fastest renewing organ provides such a system: Consisting of crypts containing intestinal stem cells (ISCs), early progenitor cells, and niche-providing Paneth cells, and villi composed of mature cell types, a spatial differentiation hierarchy guides the directional movement of ISCs during differentiation and maturation^5,6^ (Fig. 1A left). This feature is recapitulated in *in vitro* tissue models of the gut: In budding structures of 3D mouse small intestinal organoids (mSIOs; enteroids) ISC-containing domains are located at the tips and differentiating ISCs move inward^7,8^ (Fig. 1A middle). In 2D enteroid monolayer cultures (EMCs)^9–11^ ISCs in proliferative centers, also containing differentiated Paneth cells, move outward along a radial differentiation trajectory^9^ (Fig. 1A right). Such 3D/2D organoid models retain all intestinal epithelial cell types and are amenable to live imaging, which enabled the study of cellular behavior within multicellular tissue-resembling systems^12–16^, including live-cell TF dynamics at the cellular^17^ and single-molecule level^18^.

**Figure 1:**
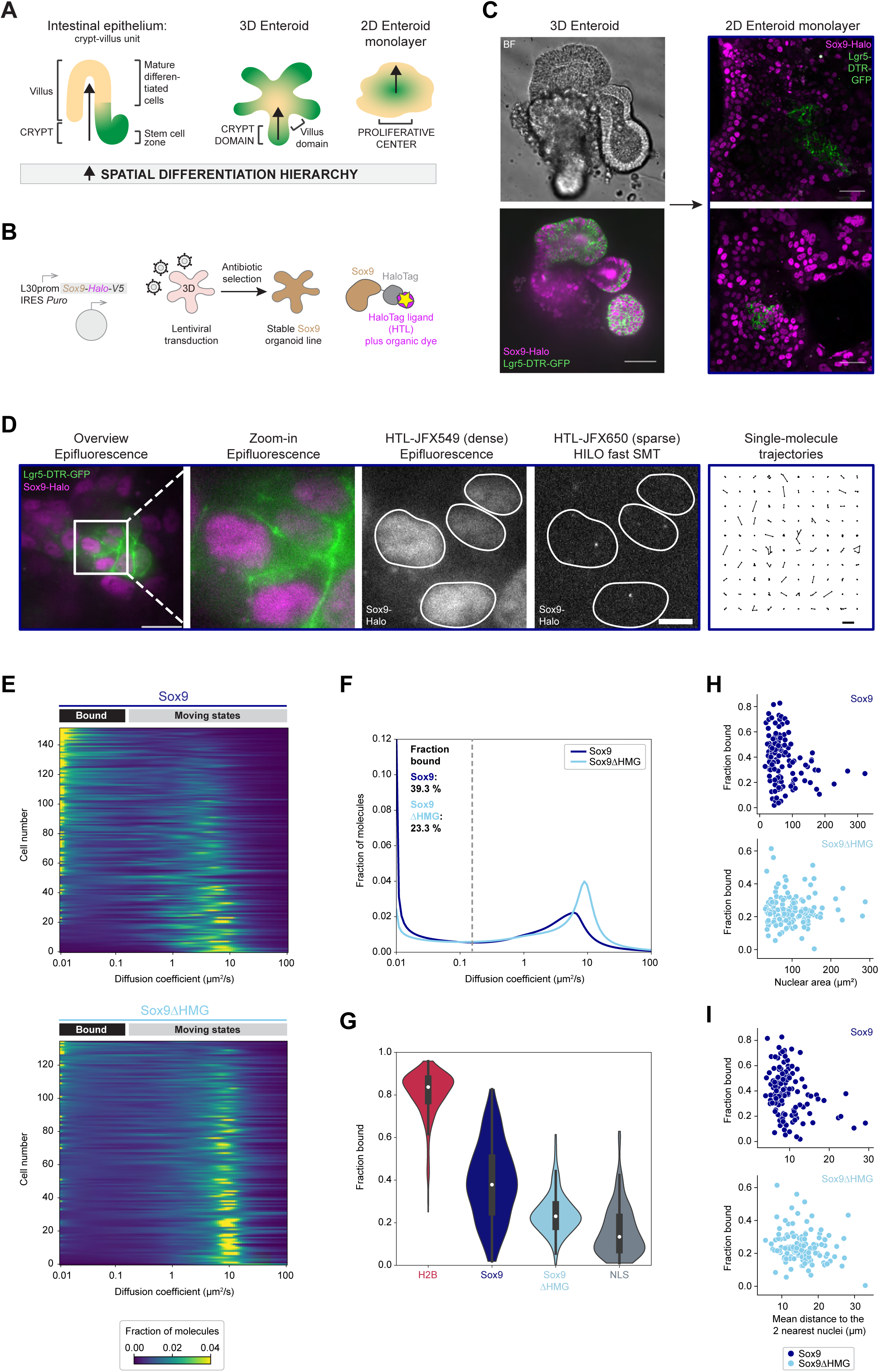
Automated live-cell fast single-molecule tracking (SMT) in 2D enteroid monolayer cultures (EMCs) reveals a heterogenous diffusive behavior of Sox9- Halo with the fraction of immobile molecules largely depending on DNA binding. **(A)** In the mammalian intestinal epithelium (left) a spatial differentiation hierarchy (arrow) guides directional movement of differentiating intestinal stem cells (ISCs) along the crypt-villus (green-beige) axis, also reflected in *in vitro* models of the intestinal epithelium: in mouse small intestinal organoids (mSIOs; enteroids; middle), ISCs in the crypt domain at bud tips (green) migrate inward upon differentiation, whereas 2D EMCs (right) grow outward from ISC-containing proliferative centers (green). **(B)** Generation of an organoid line stably expressing a Sox9-HaloTag(Halo)- V5 transgene through lentiviral transduction and antibiotic selection allows fluorescence detection (yellow star) of Sox9-Halo covalently labeled with dye-coupled HaloTag ligands (HTLs). **(C)** Confocal imaging of mSIOs (left) derived from Lgr5::DTR- GFP mice (labeling ISCs/early progenitors (green)) stably expressing Sox9-Halo (magenta) and corresponding 2D EMCs (right) 5 d post-seeding. BF: brightfield. Scale bars: 50 μm. **(D)** Double labeling of Sox9-Halo 2D EMCs with two different HTLs allows bulk labeling for nuclear segmentation (images 1-3 from left; white masks) and sparse labeling for HILO-based fast SMT (image 4; one representative frame of a SMT movie) resulting in single-molecule trajectories (image 5; 100 randomly selected single- molecule trajectories). Scale bars: 20 μm (overview), 5 μm (zoom-in), 1 μm (trajectories). **(E)** Single-cell diffusion heatmaps for 4 combined automated SMT experiments for Sox9-Halo (dark blue; *n*=152 cells) and 5 combined manual fast SMT experiments for Sox9ΔHMG-Halo (light blue; *n*=135 cells). Cells are ordered by decreasing fraction bound (top-bottom). **(F)** Mean diffusion spectra for Sox9-Halo (dark blue) and Sox9ΔHMG-Halo (light blue). Bootstrap analysis of combined experiments with *n*=10,44,42,56 cells for Sox9 and *n*=43,23,1,34,34 cells for Sox9ΔHMG determined a mean fraction bound (*D*<0.15μm^2^/s) of 39.3% (95% confidence interval (CI): 32.9-46.7%) and 23.3% (95% CI: 19.5-28.5%), respectively. **(G)** Violin plots for fractions bound (white point: median; whiskers: first/third quartile) of Halo-tagged H2B (red), Sox9 (dark blue), Sox9ΔHMG (light blue), and nuclear localization sequence (NLS; gray). Sox9 and Sox9ΔHMG data are the same as in (E,F). Data are from 7 H2B and 3 NLS experiments with combined *n*=367 and *n*=100 cells (*n*=65,30,12,50,123,81,6 and *n*=12,47,41 cells per experiment). **(H,I)** Single-cell correlation of SMT-derived fraction bound for Sox9 (top, dark blue) or Sox9ΔHMG (bottom, light blue) with the morphological characteristics nuclear area (H) or mean distance to the two nearest nuclei (I).

TFs diffuse in the nucleus in search of their cognate DNA binding motifs, to which they eventually bind to regulate target gene expression^2^. Diffusion and binding properties of single TF molecules can be measured in live cells by single-molecule tracking (SMT). In contrast to other bulk fluorescence imaging methods^19^, SMT directly allows the detection, localization and tracking of individual molecules, enabling the resolution of subpopulations with different diffusive behaviors within the same cell^20^. The obtained diffusion spectra provide the following parameters: 1) the diffusion coefficients of differentially mobile subpopulations (e.g. freely, fast, and slowly diffusing) and 2) the fraction of immobile molecules, which, in the case of TFs, are mainly chromatin-bound (fraction bound)^20,21^.

Highly inclined and laminated optical sheet (HILO)-based SMT has been used extensively to probe TF dynamics in 2D cell culture systems^22–30^, including cancer cells^22,25,29,30^, embryonic stem cells^23,26,27^, primary neurons^28^, and in distinct cell states obtained through directed differentiation protocols^24,31^, but not yet cell type-resolved within a multicellular differentiation context. 2D EMCs as multicellular differentiation systems are very heterogenous since comprised of various cell types. To deal with such complexity, we recently developed an automated SMT imaging and analysis pipeline in 2D EMCs that records hundreds of cells and resolves their heterogeneity in TF diffusion correlated with cellular features indicative of differentiation states^18^. Additionally, we implemented proximity-assisted photoactivation (PAPA)-SMT in 2D EMCs^18^, a technique probing molecular interactions in live cells at single-molecule resolution^32^ and thus resolving diffusion parameters of specific molecular complexes^18,20,29,33^.

The TF sex-determining region Y (SRY) box 9 (Sox9) exerts important functions in multiple organs^34–41^, including the intestine^42^, both during embryonic development and adult tissue differentiation. Sox9 mutations and an aberrant Sox9 dosage are implicated in a wide range of diseases, ranging from skeletal dysplasia and sex reversal^43^ to colorectal cancer^44,45^.

Sox9 contains a high mobility group (HMG) box DNA binding domain^46^ and can homodimerize via its dimerization domain (DIM). Dimerization is required for DNA binding and transactivation of cartilage-specific genes^47,48^ but it is dispensable in other contexts^48^, where Sox9 functions as monomer.

During mouse intestinal development, Sox9 is expressed in all epithelial cells at E13.5 but it becomes restricted to proliferating progenitor cells at E15.5^42^. Upon adulthood, Sox9 is expressed in ISCs, progenitor, and secretory Paneth cells^42^. Here, Sox9 is required for progenitor cell maintenance^51^ and Paneth cell differentiation^42,52^, whereby distinct Sox9 expression levels characterize various cell populations within the murine small intestinal crypt^53^.

Due to its diverse roles and expression levels during intestinal differentiation and beyond, we chose Sox9 to study how abundance and diffusive behavior of a cell fate- conferring TF change during differentiation. Using our automated SMT pipeline^18^, we investigated the dynamics of Sox9 under homeostatic conditions in differentiating 2D EMCs. By directly recording cell type markers, we determined an expression level-independent correlation between the fraction of immobile Sox9 molecules and the progression of differentiation, largely dependent on DNA binding. We further observed that long-term overexpression of Sox9 in mSIOs causes a change in organoid morphology, accompanied by increased proliferation, as well as a loss of intestinal gene expression signatures and the acquisition of a regenerative fetal-like gene expression program. Applying our (PAPA)-SMT pipelines^18^ to 3D spheroids and spheroid-derived 2D EMCs, we observed increased DNA occupancy by Sox9 and evidence of oligomerization. Our results suggest context-dependent molecular dynamics of Sox9 during differentiation in adult intestinal homeostasis and upon Sox9 overexpression-induced fetal-like reprogramming.

## Results

### Automated fast SMT in live 2D EMCs reveals heterogenous Sox9-Halo diffusion across differentiating cells independent of expression levels

To use Sox9 for interrogating the diffusive behavior of a cell fate-conferring TF in the intestinal differentiation paradigm (Fig. 1A), we used lentiviral transgene delivery (Fig. 1B) to generate a stable mSIO line expressing Sox9-Halo from a weak ubiquitous L30 promoter^22^. Nuclear Sox9-Halo expression in 3D mSIOs and 2D EMCs derived thereof was confirmed by live imaging (Fig. 1C) after covalent labeling of the HaloTag (Halo) with HaloTag ligands (HTLs) coupled to bright and photostable dyes^54^, which was also key for SMT (Fig. 1D). Here, we used HILO-based live-cell fast SMT on a total internal reflection fluorescence (TIRF) microscope and employed a stroboscopic illumination scheme to reduce motion blur^20,26^. A typical manual fast SMT experiment of 10 randomly selected cells did not provide a conclusive picture about the Sox9-Halo diffusion behavior in these heterogenous EMCs (Fig. S1A,B), demonstrating the need to acquire larger datasets. We thus used our previously developed automated SMT pipeline^18^ to acquire hundreds of cells (Fig. 1D; Fig. 1E top; Fig. S1E). Bulk-labeled Sox9-Halo (Fig. 1D, subpanels 1-3) allowed the automated detection of nuclei and triggered an SMT sequence in another channel with Sox9-Halo sparsely labeled with a second HTL-coupled fluorophore (Fig. 1D, subpanel 4) to obtain single-molecule trajectories^18^ (Fig. 1D, subpanel 5). Automated SMT in hundreds of randomly chosen cells confirmed a large heterogeneity in the diffusive behavior of Sox9-Halo with diffusion peaks ranging from freely diffusing (*D* ∼10 µm^2^/s) to immobile (*D* ∼0.01 µm^2^/s) (Fig. 1E, top; Fig. S1E) and an average fraction of immobile (*D* <0.15 µm^2^/s) Sox9- Halo molecules of 39.3% (95% confidence interval (CI): 32.9–46.7% Fig. 1F,G). Notably, cell-to-cell differences in the fraction bound covered the whole spectrum between the average fractions bound determined for the immobile H2B-Halo (83.7%; 95% CI: 81.4–85.9%) and freely diffusing Halo-nuclear localization sequence (NLS; 19.2%; 95% CI: 15.0–23.2%) controls (CTRLs) (Fig. 1G; Fig. S1I). On a cell-by-cell basis, the fractions bound of Sox9-Halo did not correlate with the average nuclear Sox9-Halo intensity (Fig. S1G), a proxy for transgenic Sox9-Halo levels, arguing against heterogeneity being caused by variable transgene expression in the initially polyclonal organoid line.

### Immobile Sox9-Halo molecules largely reflect DNA binding

A plausible explanation for the observation of immobile states of TFs in the cell nucleus is TF binding to DNA. To test whether this is the case for the immobile Sox9-Halo molecules detected by SMT, we transiently expressed a Sox9-Halo mutant lacking the HMG DNA binding domain (Sox9ΔHMG-Halo) in wild-type (WT) organoid-derived 2D EMCs via recombinant adeno-associated viral (rAAV) vector delivery^55^ (Fig. S1C,D). Automated fast SMT of Sox9ΔHMG-Halo revealed a more uniform diffusive behavior (Fig. 1E bottom; Fig. 1G; Fig. S1F) with a lower average fraction bound of 23.3% (95% CI: 19.5–28.5%) compared to full-length Sox9-Halo (Fig. 1F,G) independent of its nuclear expression level (Fig. S1H). Hereby, most cells were characterized by a diffusion peak in the freely diffusing range, confirming that the measured immobile fraction largely represents Sox9-Halo molecules bound to DNA. Nevertheless, the average fraction bound of Sox9ΔHMG-Halo did not fully decrease to the 19.2% determined for the Halo-NLS freely diffusing CTRL (Fig. 1G; Fig. S1I), suggesting that additional Sox9 protein domains might contribute to the bound fraction measured by SMT.

### Cells with more immobile Sox9-Halo molecules display morphological features of intestinal stem/early progenitor cells in 2D EMCs

Hypothesizing that the observed cell-to-cell heterogeneity in the diffusive behavior of Sox9-Halo might reflect differentiation states, we inspected our single-cell diffusion data with respect to nuclear area and the mean distance of a nucleus to its two nearest nuclei, two parameters extracted from our images that we previously found to be correlated with differentiation^18^. For Sox9-Halo, we indeed identified a subpopulation of cells with smaller nuclei or a smaller nearest-nuclei-distance, indicative of stem/early progenitor cells in proliferative centers^18^ (Fig. 1C right), which was characterized by larger fractions bound (Fig. 1H top). In contrast, a subpopulation of cells with larger nuclei or a larger nearest-nuclei-distance, indicative of differentiated cells further away from proliferative centers^18^ (Fig. 1C right), was characterized by a smaller fraction bound (Fig. 1I top). These subpopulations with distinct diffusive and morphological features were not discernible for the Sox9ΔHMG-Halo mutant (Fig. 1H,I), suggesting that the immobile fraction of Sox9-Halo in stem/early progenitor cells mainly reflects DNA binding.

### DNA-binding of Sox9-Halo decreases upon intestinal differentiation

To directly test our observation of distinct diffusive subpopulations with morphological characteristics indicative of differentiation states, we implemented the recording of fluorescent cell type markers and marker-based cell classification into categories in our SMT pipeline^18^. Using the green fluorescent ISC/early progenitor marker Lgr5, present in our Sox9-Halo organoid line derived from Lgr5::DTR-GFP mice^56^, we classified the Sox9-Halo population into Lgr5-positive (Lgr5+) stem/early progenitor cells and Lgr5-negative (Lgr5-) late progenitor/differentiated cells (Fig. 2A). The average fraction bound of Sox9-Halo decreased during differentiation from 48.2% (95% CI: 44.2–52.3%) in stem/early progenitor cells to 38.1% (95% CI: 31.9–44.5%) in late progenitor/differentiated cells (Fig. 2B; Fig. 2C right; Fig. S2A,B; Fig. S2C bottom). Such decrease was also apparent upon hierarchical clustering of Lgr5+/- single-cell diffusion spectra together^18^ (Fig. S2D-G). Notably, cell-to-cell variability persisted within both Lgr5+/- subpopulations (Fig. 2C right; Fig. S2A,B; Fig. S2C bottom), indicating a potentially more complex grading of the Sox9 diffusion behavior into cell states and types within these heterogenous subpopulations. As observed for the whole population (Fig. S1G), the differences in the diffusive behavior of Sox9-Halo were independent of its expression level in both Lgr5+/- subpopulations (Fig. 2D). Importantly, for both H2B-Halo and Halo-NLS immobile and freely diffusing CTRLs, respectively, no difference in the diffusive behavior was measured between Lgr5+/- subpopulations (Fig. S2H,I), further supporting that the observed difference in the Sox9-Halo diffusion behavior stems from differentiation and not from other differences between stem and differentiated cells, such as nuclear size or crowding. Notably, the Lgr5+ subpopulation was characterized by smaller nuclei and a smaller nearest- nuclei-distance in comparison to the Lgr5- subpopulation (Fig. 2C left; Fig. S2C top), further validating our cell morphology-based approach for an approximated discrimination of undifferentiated from differentiated cells in 2D EMCs^18^ (Fig. 1H,I). Nevertheless, we noted some Lgr5- cells with smaller nuclei and larger fractions bound (Fig. 2C,D), for which one possible explanation could be differentiated Paneth cells residing within proliferative centers. Taken together, consistent with Sox9 functioning in the proliferative intestinal crypt, our results suggest that the Lgr5+ subpopulation is characterized by more Sox9-Halo molecules bound to DNA, possibly resulting from the occupancy of more DNA binding sites.

**Figure 2:**
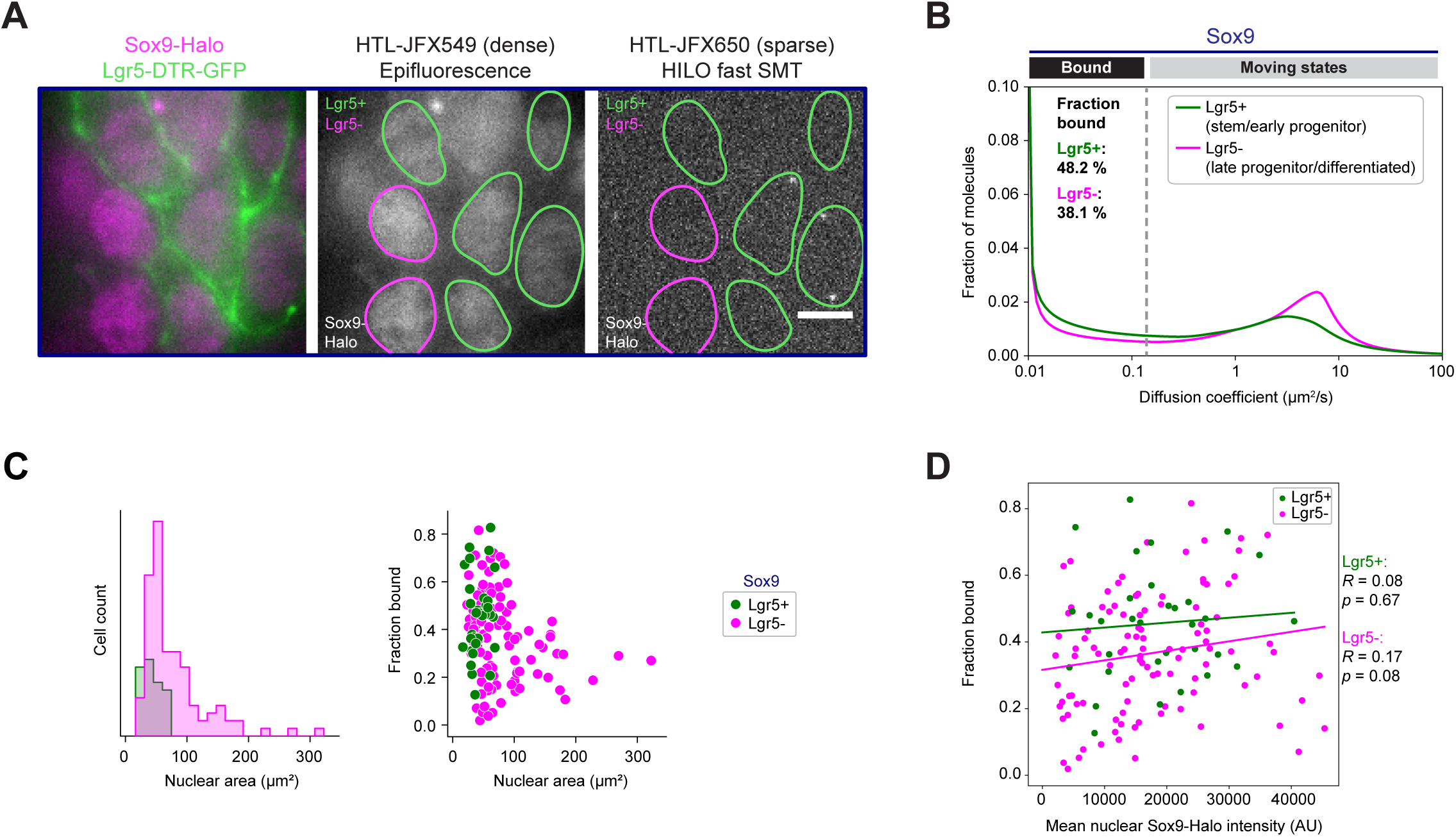
Lgr5-based distinction of differentiation states demonstrates an expression level-independent decrease in the fraction of immobile Sox9-Halo molecules upon differentiation. **(A)** The green fluorescent Lgr5-DTR-GFP marker enables classification into stem/early progenitor (Lgr5+) and late progenitor/differentiated cells (Lgr5-) for SMT in 2D EMCs. Left: epifluorescence of bulk-labeled Sox9-Halo (magenta) and Lgr5-DTR-GFP (green); middle: epifluorescence of bulk-labeled Sox9-Halo (gray) with segmented nuclear masks color-coded according to classification into Lgr5+/- (green/magenta) categories; right: representative frame of a fast SMT movie of sparsely labeled Sox9-Halo (gray) with same classified masks depicted. Scale bar: 5 μm. **(B)** Mean diffusion spectra of Sox9- Halo for Lgr5+/- (green/magenta) subpopulations. Bootstrap analysis of 4 combined experiments with *n*=3,21,2,10 Lgr5+ and *n*=7,23,40,46 Lgr5- cells determined a mean fraction bound of 48.2% (95% CI: 44.2-52.3%) and 38.1% (95% CI: 31.9-44.5%), respectively. **(C)** Left: Nuclear area distribution of Lgr5+/- (green/magenta) cells extracted from SMT data. Right: Single-cell correlation of SMT-derived fraction bound with nuclear area for Lgr5+/- cells. **(D)** Fractions bound for each cell plotted against the mean nuclear Sox9-Halo intensity for Lgr5+/- cells. Correlations (fitted lines) were computed for each subpopulation; Pearson correlation coefficients (*R*) and *p*-values are indicated. The data in (B-D) are the same as in Fig. 1E-I, Fig. 5C, Fig. S1E,G,I, and Fig. S2A-G.

### Long-term overexpression of Sox9-Halo results in a proliferative cell state transition accompanied by an organoid morphology change to spheroids devoid of signatures for intestinal identity and function

Prolonged culture of a stable Sox9-Halo organoid line resulted in progressive changes in organoid morphology: starting from the initially typical budding morphology (until ∼10 weeks after line establishment (LE)) organoids got rounder with smaller buds (∼15-25 weeks after LE) and finally completely spherical (∼30 weeks after LE) (Fig. 3A, Fig. S3A left). In contrast, non-transduced WT organoids continued to grow as budding structures until our longest observed culture time of ∼22 weeks (Fig. 3A). This morphological change in 3D Sox9-Halo organoids was also reflected in 2D EMCs derived from them, which were characterized by smoother edges of the monolayer in comparison to WT EMCs (Fig. 3A) and grew to complete confluency (Fig. S3A right), suggesting a loss in cell contact inhibition. While Sox9 expression was no longer restricted to proliferative centers^42^, as expected for ubiquitous Sox9-Halo transgene expression, the average expression level of total (endogenous plus transgene) Sox9 protein in spheroids was only ∼2.3x higher than the endogenous Sox9 level in proliferative centers of WT organoids (Fig. 3C left), while the average Sox9-Halo transgene expression level was similar to that in budding Sox9-Halo organoids (Fig. 3C right). Notably, Sox9 spheroids were characterized by on average larger nuclei (Fig. 3D left) and a larger nearest-nuclei-distance (Fig. 3D right) than WT organoids, indicative of a more differentiated cell population^18^ (Fig. 2C; Fig. S2C). Despite such morphology, spheroids contained more (76.4%) proliferative cells than WT organoids (51.1%) as determined by immunofluorescence (IF) for the proliferation marker Ki67^57^ (Fig. 3B,E), consistent with the described role for Sox9 in maintaining proliferative progenitor cells^51^. This increased proliferative capacity was accompanied by several aberrant cell division phenotypes, e.g. fusion of two or more nuclei, micronuclei and fragmented nuclei (Fig. S3B).

**Figure 3:**
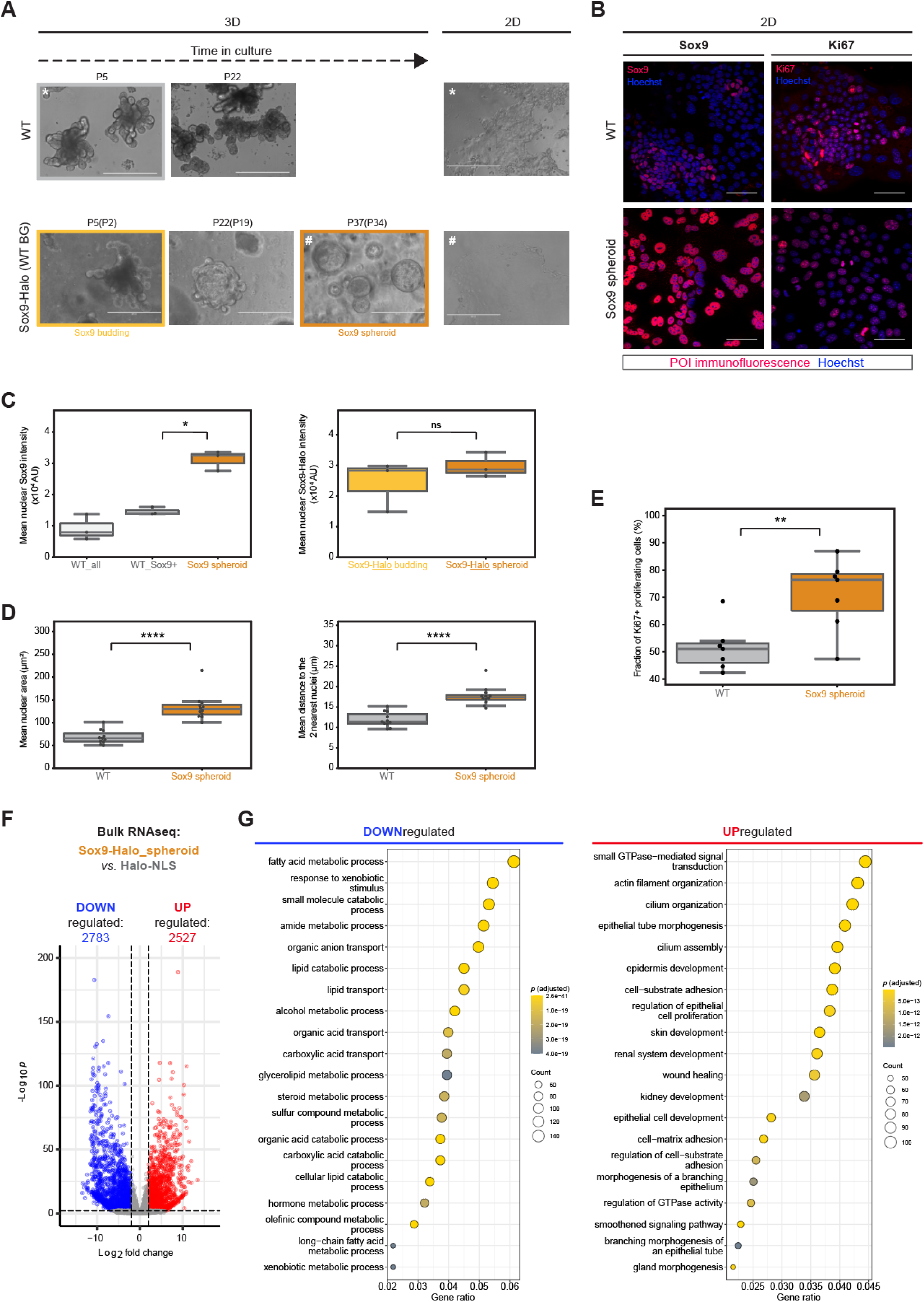
Long-term overexpression of Sox9-Halo in enteroids results in an organoid morphology change coinciding with a proliferative cell state transition and a loss of gene expression signatures for intestinal identity and function. **(A)** In contrast to wild-type (WT) organoids (top), stable Sox9-Halo organoids (WT background; bottom) transitioned from budding to spheroid morphology upon long- term culture. This morphology change of 3D organoids (left) is also reflected in 2D EMCs derived from them (right). Epifluorescence images 5 d post-seeding with organoid passage number P(P) after crypt isolation (after lentiviral transduction to make stable line) indicated. */# denote passage of 2D EMC derivation from 3D organoids. Scale bars: 400 μm. **(B)** Confocal images of 2D EMCs derived from WT organoids (top) or Sox9-Halo spheroids (bottom) immunostained (red) for Sox9 (left) or the proliferation marker Ki67 (right) and co-stained with Hoechst (blue) 5 d post- seeding. Scale bars: 50 μm. **(C)** Quantification of total (left) or transgene (right) Sox9 protein levels in 2D EMCs derived from WT (gray), Sox9-Halo_budding (yellow) or Sox9-Halo_spheroid (orange) organoids based on fixed and immunostained (left) or live and HTL-stained (right) confocal images 5 d post-seeding. Total Sox9 levels: 3 FOVs with 456 cells (162 Sox9+, 294 Sox9-) (WT), 3 FOVs with 159 cells (Sox9_spheroid). Transgene Sox9-Halo levels: 3 FOVs with 864 cells (Sox9_budding), 3 FOVs with 537 cells (Sox9_spheroid). **(D)** Quantification of nuclear area (left) and mean distance to the two nearest nuclei (right) in 2D EMCs derived from WT organoids (gray) or Sox9-Halo spheroids (orange) 5 d post-seeding based on confocal images (WT: 23 FOVs, 3029 cells; Sox9_spheroid: 30 FOVs, 3100 cells). **(E)** Percentage of Ki67+ proliferative cells in 2D EMCs derived from WT organoids (gray) or Sox9 spheroids (orange) quantified from confocal images as in (B). WT: 7 FOVs, 752 cells; Sox9_spheroid: 7 FOVs, 648 cells. For (C-E): Each point represents one FOV; median: gray line, first/third quartile: whiskers; statistical testing based on Mann-Whitney U tests (see methods for details); (ns) non-significant, *p*>0.5; (*) *p*≤0.5; (**) *p*≤0.01; (****) *p*≤0.0001. **(F,G)** Bulk RNAseq experiment of Sox9-Halo spheroids *versus* Halo-NLS control (CTRL) organoids in biological triplicates. (F) Volcano plot displaying differentially expressed genes (DEGs; adjusted *p*-value ≤0.01, fold change ≥2 and mean counts ≥10; red/blue: up-/downregulated). (G) Gene ontology (GO) analysis of the top 20 biological pathways enriched in DEGs down- (left, blue) or upregulated (right, red) with adjusted *p*-values and gene counts indicated. Bulk RNAseq data are the same as for Halo-NLS and Sox9-Halo spheroid samples in Fig. 4B-D and Fig. S4,5.

To reconcile the seemingly opposing effects of long-term Sox9-Halo overexpression, we performed bulk RNAseq of Sox9-Halo organoids in comparison to CTRL organoids stably expressing Halo-NLS, revealing thousands of differentially expressed genes (DEGs) (Fig. 3F). Gene ontology (GO) analysis of the 2783 genes downregulated in spheroids revealed the top 20 biological processes to be metabolic and metabolite transport processes (Fig. 3G left), pointing towards a loss of the intestinal epithelial function in nutrient absorption and metabolism. Conversely, GO analysis of the 2527 upregulated genes revealed pronounced association with epithelial morphogenesis and the development and differentiation of various non-intestinal tissues and organs (Fig. 3G right), indicating a loss in intestinal identity and the activation of a gene expression program resembling embryonic-like development. Wound healing, cell adhesion, actin filament organization, and cilium assembly were among other biological processes associated with genes upregulated in spheroids (Fig. 3G right), some of which we phenotypically confirmed by IF, including the formation of actin stress fibers and membrane spikes (Fig. S3C). These results altogether suggest a large functional and epithelial rearrangement underlying Sox9-Halo spheroid formation.

### Sox9 overexpression results in a transient upregulation of stem cell markers followed by Yap activation and a regenerative fetal-like gene expression program

Given the extensive gene expression rearrangement in Sox9 spheroids, we questioned whether the HaloTag fused to the Sox9 transgene might have contributed to its emergence. Reassuringly, the budding-to-spheroid morphology transition was reproduced in a stable Sox9-mEGFP organoid line (Fig. 4A) on a similar time scale, ruling out potential effects due to the introduced HaloTag. To further compare the two stable Sox9 organoid lines and gain insights into the gene expression changes underlying organoid morphology transition, we selected one budding and one spheroid phenotype for each line and ordered them phenotypically based on the progression of spheroid morphology acquisition, which also correlated with their time in culture (Fig. 4A). The addition of three intermediate phenotypes (Sox9-mEGFP_budding, Sox9- Halo_budding, Sox9-mEGFP_spheroid) (Fig. 4A) to our previously analyzed Sox9- Halo spheroid and Halo-NLS organoid samples (Fig. 3F,G) allowed us to perform a morphology “pseudo-time-course analysis” (Fig. 4A). Both spheroid samples were very different from the two budding and the Halo-NLS CTRL samples (Fig. S4A), which clustered together upon principal component analysis (PCA) (Fig. 4B). Nevertheless, comparing Sox9-mEGFP *versus* Sox9-Halo budding and spheroid samples, about a hundred (Fig. S4B) or one thousand DEGs (Fig. S4C) were detected, respectively, confirming that gene expression changes underlie the slightly different morphology phenotypes and ultimately lead to Sox9 spheroid formation (Fig. 3F, Fig. 4A).

**Figure 4:**
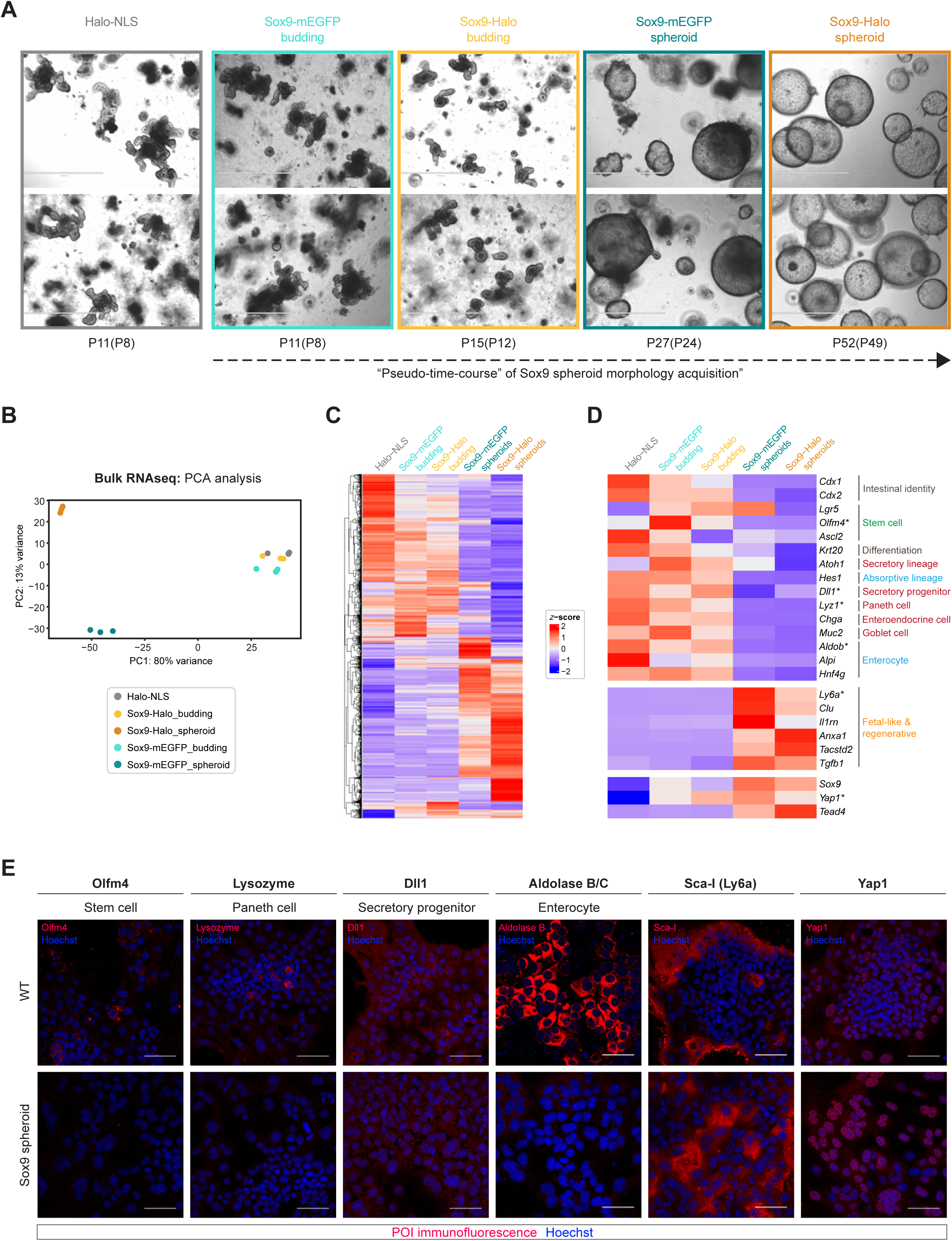
A “pseudo-time-course” of spheroid morphology acquisition upon Sox9 overexpression reveals a transient increase in stem cell markers followed by a reduction in intestinal epithelial signatures towards a fetal-like reversion. **(A)** Representative images of the stable organoid lines Halo-NLS (gray), Sox9- mEGFP_budding (light turquoise), Sox9-Halo_budding (yellow), Sox9- mEGFP_spheroid (dark turquoise), and Sox9-Halo_spheroid (orange) at the indicated passage numbers P(P) after crypt isolation (after lentiviral transduction to make stable lines) 5 d post-seeding, chronologically ordered. Scale bars: 1 mm. **(B)** Principal component analysis (PCA) of bulk RNAseq experiments of the samples (biological triplicates) in (A). **(C,D)** *z-*score heatmaps of all DEGs (C) or selected genes (D) determined by bulk RNAseq (adjusted *p*-value ≤0.01, fold change ≥2 and mean counts ≥10; red/blue: up-/downregulated). **(E)** Confocal images of immunostained (POI: protein of interest) 2D EMCs derived from WT organoids (top) or Sox9-Halo spheroids (bottom) co-stained with Hoechst (blue) 5 d post-seeding. Scale bars: 50 μm. RNAseq data in (B-D) are the same as in Fig. S4,5. RNAseq data from Halo-NLS and Sox9- Halo_spheroid samples are the same as in Fig. 3F,G.

Focusing on selected markers of intestinal epithelial lineages, we confirmed our initial GO analysis results (Fig. 3G) with a loss of intestinal identity genes, stem cell markers, and markers of absorptive and secretory lineages^58^ (Fig. 4D; Fig. S5A), some of which we confirmed at the protein level by IF (Fig. 4E). Inspecting more carefully the three samples Halo-NLS, Sox9-Halo_budding and Sox9-Halo_spheroid revealed an initial mild gene expression deregulation (Fig. S4D) followed by a second wave of major entity (Fig. S4F). While both of these transitions were characterized by an overall downregulation of metabolic and metabolite transport processes (Fig. S4E,G left), including digestion for the second transition (Fig. S4G left), genes upregulated in the first transition were associated with the Wnt signaling pathway and its regulation (Fig. S4E right), followed by upregulation of genes involved in development and differentiation of various non-intestinal tissues and organs in the second transition (Fig. S4G right). This transient upregulation of genes associated with Wnt signaling was in agreement with a transient increase in canonical stem cell markers including *Olfm4*^59^ and the Wnt target gene *Lgr5*^60^ in budding Sox9 organoids (Fig. 4D; Fig. S5A). Instead of intestinal epithelial genes, spheroids were characterized by the expression of several genes previously associated with fetal-like and regenerative signatures (e.g. *Ly6a*, *Clu*, *Anxa1*) (Fig. 4D; Fig. S5B; Fig. 4E) typically arising upon tissue regeneration after exposure of the intestinal epithelium to various sources of damage^61^.

The effector of the Hippo signaling pathway, Yes-associated protein (Yap1), has been associated with intestinal regeneration^62–65^. While mRNA levels of *Yap1* were not significantly altered in the course of Sox9 spheroid phenotype acquisition (Fig. S5C), 2D EMCs derived from these spheroids were characterized by an overall nuclear localization of Yap1 (Fig. 4E), indicative of Yap1 activation^66^. Furthermore, inspection of our bulk RNAseq data revealed an increased expression of *Tead4* (Fig. 4D; Fig. S5C), encoding a TEA domain (TEAD) TF interacting with nuclear Yap1 and thereby activating downstream genes^67^.

Taken together, our “pseudo-time-course” of spheroid morphology acquisition upon Sox9 overexpression revealed that a loss of adult homeostatic intestinal identity signatures was counteracted by a gain in fetal-like regenerative signatures. Hereby, a transient activation of Wnt signaling seems to be accompanied by a persistent activation of Yap1 signaling, resulting in a regenerative fetal-like reprogrammed phenotype.

### Increased DNA-binding of Sox9-Halo in Sox9 spheroid-derived EMCs in an expression level-dependent manner

To investigate whether the diffusive behavior of Sox9-Halo changed during fetal-like reversion, we performed automated fast SMT in 2D EMCs derived from Sox9-Halo spheroids (Fig. 5A). Here, the diffusive behavior of Sox9-Halo was more uniform with a dominating immobile diffusion peak for most of the about 500 cells measured (Fig. 5B; Fig. S6A). Indeed, Sox9-Halo in 2D EMCs derived from spheroids was characterized by a higher fraction bound of 61.1% (95% CI: 55.4–66.1%) in comparison to 39.3% in those derived from budding organoids (Fig. 5C; Fig. S6A,B), raising the possibility that more Sox9 molecules bound to DNA served as molecular driver of the observed proliferative cell state transition underlying spheroid formation. This increase in fraction bound was also apparent upon hierarchical clustering of the fast SMT data (Fig. S6C-F). Notably, unlike the budding state, in the spheroid state we measured a positive correlation between Sox9-Halo expression levels and fraction of immobile molecules (Fig. 5D), indicating that despite the presence of more Sox9-Halo molecules (Fig. 3C) a larger fraction of them was immobile in the cell nucleus and thus presumably DNA-bound.

**Figure 5:**
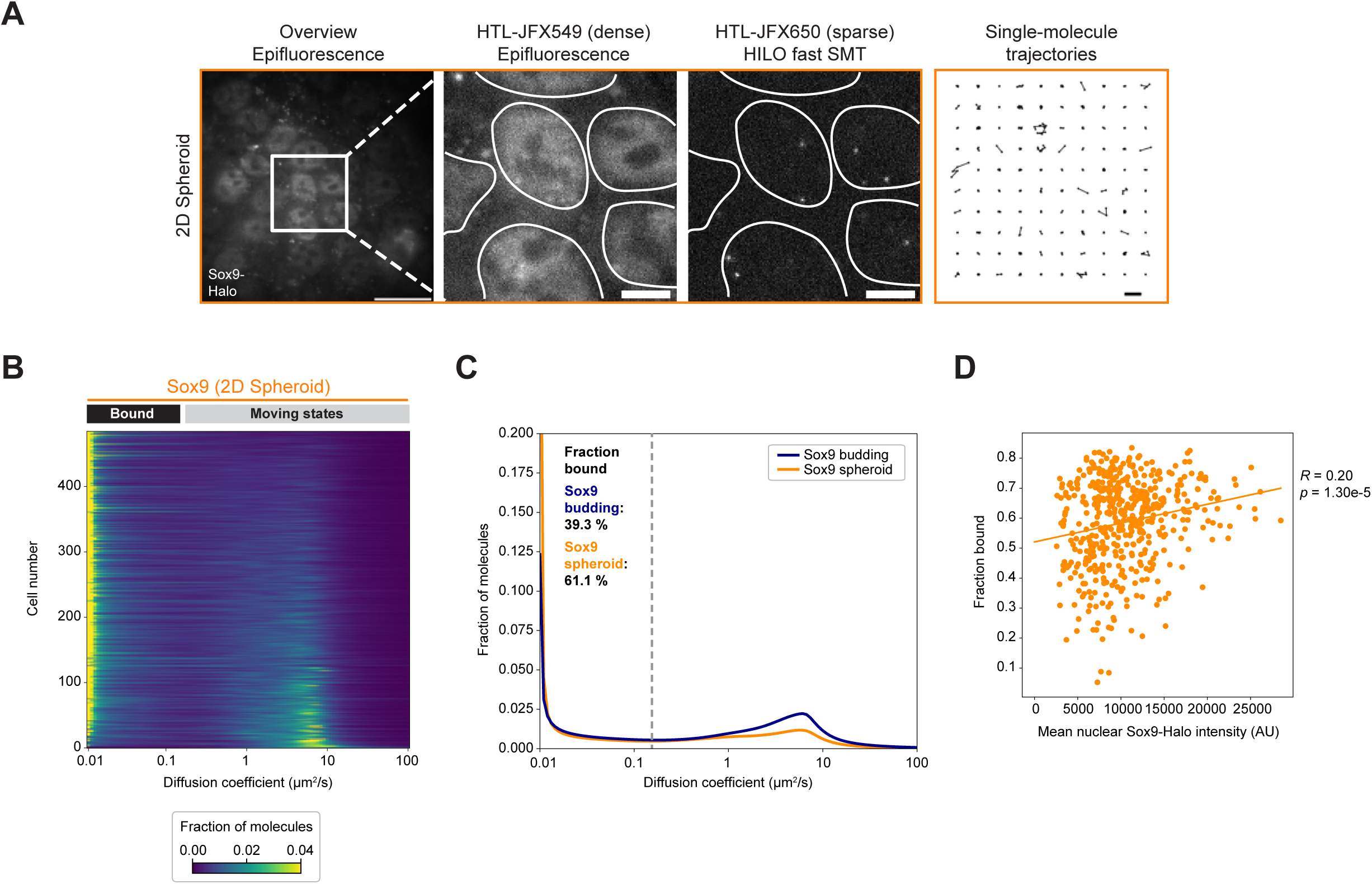
Automated fast SMT in Sox9 spheroid-derived EMCs reveals an expression level-dependent increase in the fraction of immobile Sox9-Halo molecules. **(A)** Double labeling of Sox9-Halo (gray) with two different HTLs allows bulk labeling (images 1,2 from left) for nuclear segmentation (images 2,3; white masks) and sparse labeling for fast SMT (image 3; one representative frame of a SMT movie) resulting in single-molecule trajectories (image 4; 100 randomly selected single-molecule trajectories). Scale bars: 20 μm (overview), 5 μm (zoom-in), 1 μm (trajectories). **(B)** Single-cell diffusion heatmap for 10 combined automated SMT experiments for Sox9-Halo in spheroid-derived 2D EMCs (*n*=485 cells with 77,73,13,24,30,58,81,71,26,32 cells per experiment). Cells are ordered by decreasing fraction bound (top to bottom). **(C)** Mean diffusion spectra for Sox9-Halo in 2D EMCs derived from budding organoids (Lgr5::DTR-GFP background; dark blue) or spheroids (WT background; orange). Bootstrap analysis of combined experiments with *n*=10,44,42,56 cells for Sox9-Halo_budding and *n*=77,73,13,24,30,58,81,71,26,32 cells for Sox9-Halo_spheroid determined a mean fraction bound of 39.3% (95% CI: 32.9-46.7%) and 61.1% (95% CI: 55.4-66.1%), respectively. **(D)** Fractions bound for each spheroid-derived cell plotted against the mean nuclear Sox9-Halo intensity. The correlation (fitted line) was computed; Pearson correlation coefficient (*R*) and *p*-value are indicated. The SMT data for Sox9_budding are the same as in Fig. 1E-I, Fig. 2B-D, Fig. S1E,G,I, and Fig. S2A-G.

### Proximity-assisted photoactivation (PAPA)-SMT reveals a chromatin-bound pool of self-associated Sox9 in live 2D EMCs

The increased fraction of Sox9 molecules bound to DNA in spheroids raised the question of whether Sox9 molecules might bind to DNA in an oligomerized state, as both Sox9 monomers and dimers have been reported as transcriptionally active units depending on the differentiation context^47,48^. To address this, we employed PAPA-SMT, a recently developed single-molecule method allowing to detect whether differentially labeled sender and receiver molecules (Fig. 6A) are associated or in vicinity to each other by using a PAPA illumination scheme (PAPA) distinct from that used in standard SMT (direct reactivation (DR)) (Fig. 6B). To this end, we transiently expressed Sox9- SNAPf (Fig. 6C) or a SNAPf-3xNLS CTRL (Fig. 6D) via rAAV-based delivery^55^ in Sox9- Halo spheroid-derived EMCs. PAPA-SMT using Sox9-Halo as a sender and Sox9- SNAPf as a receiver (Fig. 6A,B) revealed both freely diffusing and immobile oligomerized states (Fig. 6E) with a ∼11% higher bound fraction for PAPA *versus* DR trajectories (Fig. 6E,G), indicating a chromatin-bound pool of oligomerized Sox9. In contrast, no difference between the bound fractions of PAPA *versus* DR trajectories was detected for CTRL experiments with SNAPf-3xNLS as the receiver, measuring non-specific background PAPA^33^ (Fig. 6F,H). Notably, self-associated Sox9 was also found to be partially chromatin-bound in 2D EMCs derived from budding Sox9-Halo organoids (Fig. S7), arguing against DNA binding of an oligomerized form of Sox9 as a feature exclusive to spheroids. We nevertheless cannot exclude the existence of differences in the fraction of Sox9 molecules present in an oligomerized state or the number of Sox9 units that DNA-bound Sox9 oligomers consist of, as this PAPA-SMT assay does not provide information on whether more or larger Sox9-Halo oligomers might be immobile in the spheroid compared to the budding state.

**Figure 6:**
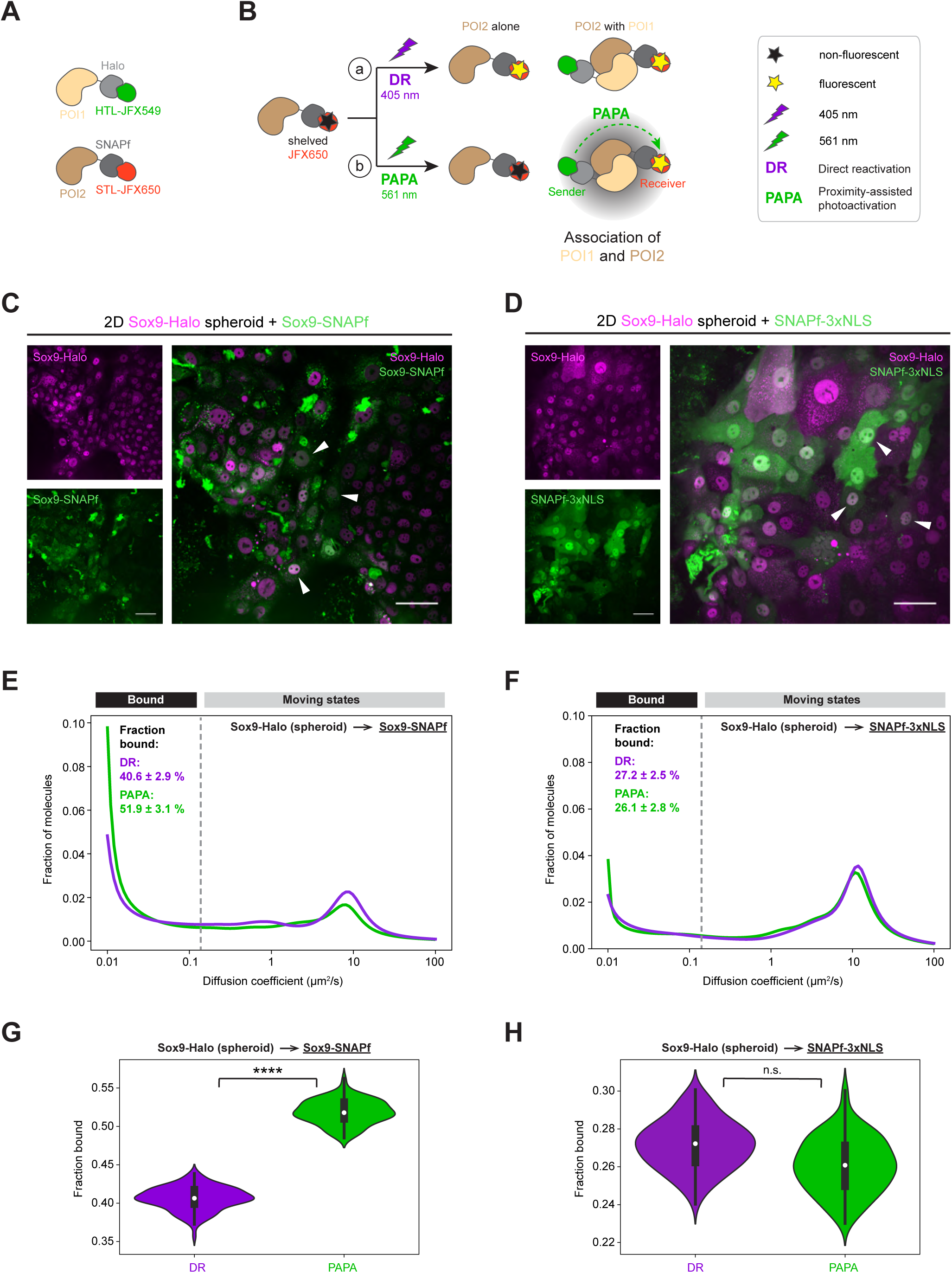
Proximity-assisted photoactivation (PAPA)-SMT reveals a chromatin-bound pool of self-associated Sox9 in spheroid-derived live 2D EMCs. **(A)** Differential labeling of POI1-Halo (beige-light gray) and POI2-SNAPf (brown-dark gray) with different HTL/SNAPfTag ligand (STL)-coupled fluorophores (JFX549- sender, green; JFX650-receiver, red) for PAPA-SMT. **(B)** Principle underlying PAPA-SMT: Through excitation using 639 nm light, shelved JFX650-labeled receiver molecules (red; black star) can either undergo a) direct reactivation (DR; purple) upon illumination with a violet 405 nm light pulse (purple lightening arrow) making them detectable independent of any potential oligomerization states as in standard SMT, or b) PAPA (green) upon illumination with a green 561 nm light pulse (green lightening arrow) making them only detectable if in proximity (gray cloud) to a JFX549-labeled sender molecule (green). **(C,D)** Confocal images of Sox9-Halo spheroid-derived 2D EMCs transduced with crude recombinant adeno-associated viral (rAAV) vectors for transient expression of (C) Sox9-SNAPf or (D) SNAPf-3xNLS (CTRL) for PAPA experiments (sender: Sox9-Halo (magenta); receiver: SNAPf-tagged component (green)). Scale bars: 50 μm. **(E,F)** Mean PAPA (green) *versus* DR (purple) diffusion spectra for PAPA-SMT of Sox9-Halo_spheroid→Sox9-SNAPf (E) or Sox9- Halo_spheroid→SNAPf-3xNLS (F) in 2D EMCs. **(G,H)** Violin plots for fractions bound (white point: median; whiskers: first/third quartile) of Sox9→Sox9 (G) and Sox9→NLS (H) determined from DR/PAPA (purple/green) trajectories. Data in (E,G) are from 4 combined experiments with *n*=129 cells (*n*=35,30,28,36 cells; 14156 DR and 9255 PAPA trajectories; bootstrapped fractions bound: 40.6±2.9% (DR), 51.9±3.1% (PAPA)). Data in (F,H) are from 3 combined experiments with *n*=70 cells (*n*=21,30,19 cells; 7781 DR and 4117 PAPA trajectories; bootstrapped fractions bound: 27.2±2.5% (DR), 26.1±2.8 (PAPA)). (n.s.) *p*>0.05, (****) *p*≤0.0001. For statistical details see methods.

### Fast SMT in 3D Sox9 spheroids confirms increased Sox9 binding upon fetal-like reversion

Finally, we sought to confirm the Sox9-Halo diffusive behavior observed in spheroid-derived 2D EMCs directly in 3D spheroids. As our HILO-based SMT approach is limited to the 10-20 µm of the sample directly above the culture glass surface, we optimized our organoid seeding procedure so that growing round spheroids could reach the glass surface, making them amenable to SMT. Using an experimental approach analogous to 2D EMCs (Fig. 7A), SMT in spheroids revealed a rather homogenous diffusive behavior of Sox9-Halo across the 35 manually imaged cells (Fig. 7B,C) with most of them being characterized by an immobile diffusion peak (Fig. 7B). With an average fraction bound of 65.6% (95% CI: 62.5–68.8%) (Fig. 7C) the diffusive behavior of Sox9-Halo in 3D spheroids is thus in agreement with that measured in spheroid-derived 2D EMCs (Fig. 5C).

**Figure 7:**
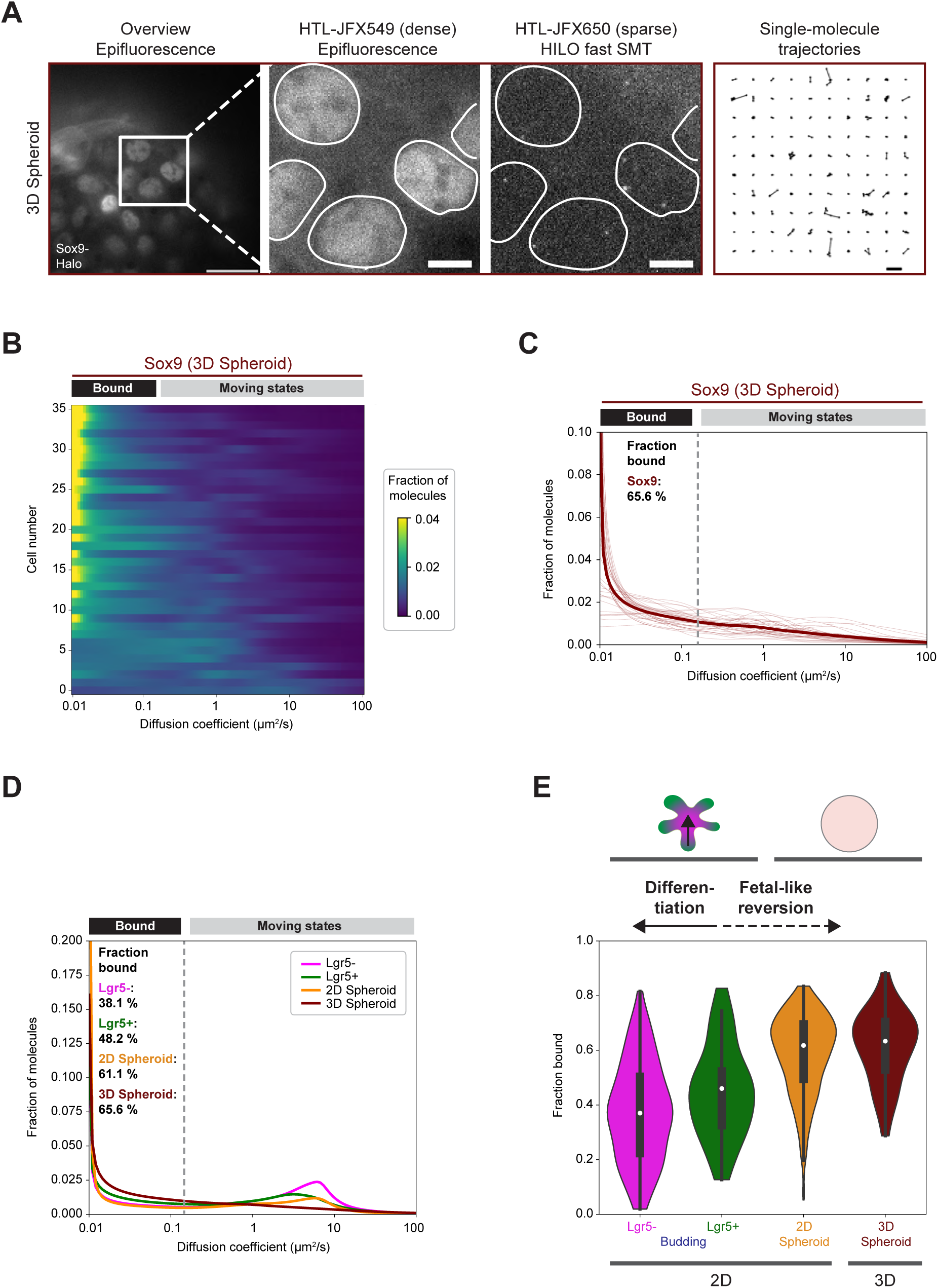
SMT in 3D spheroids confirms an increased fraction of DNA-bound Sox9-Halo molecules upon fetal-like reversion. **(A)** Manual SMT pipeline for Sox9- Halo in 3D spheroids. Double labeling of Sox9-Halo (gray) with two different HTLs allows bulk labeling (images 1,2) for nuclear segmentation (images 2,3; white masks) and sparse labeling for HILO-based fast SMT (image 3; one representative frame of a SMT movie) resulting in single-molecule trajectories (image 4; 100 randomly selected single-molecule trajectories). Scale bars: 20 μm (overview), 5 μm (zoom-ins), 1 μm (trajectories). **(B)** Single-cell diffusion heatmap for 2 combined manual SMT experiments for Sox9-Halo in 3D spheroids (combined *n*=36 cells with *n*=26,10 cells per experiment). Cells are ordered by decreasing fraction bound from top to bottom. **(C)** Single-cell diffusion spectra for data in (B). Bootstrap analysis of combined experiments with *n*=26,10 cells determined a mean fraction bound of 65.6% (95% CI: 62.5-68.8%). **(D,E)** Summary of the diffusive behavior of Sox9-Halo determined by fast SMT under homeostatic differentiation conditions (budding) and upon Sox9 overexpression-induced fetal-like reversion (spheroid). (D) Mean diffusion spectra for the Sox9-Halo_budding Lgr5+/- (green/magenta) subpopulations and for Sox9- Halo_spheroid determined in 3D spheroids (brown) or 2D EMCs (orange) with fractions bound indicated. (E) Violin plots for fractions bound (white point: median; whiskers: first/third quartile) of samples in (D).

## Discussion

In this study, we combined *in vitro* models of the rapidly renewing small intestinal epithelium (Fig. 1A) with automated live-cell SMT^18^ to investigate how the molecular dynamics and abundance of a cell fate-conferring TF change during differentiation. We further interrogated the robustness of cell fate determination to aberrant TF expression and how TF excess might alter TF molecular dynamics. Focusing on Sox9, our results show that different cellular states in adult intestinal organoid models are characterized by distinct Sox9-Halo diffusion dynamics (Fig. 7D,E): Under homeostatic conditions, the fraction of DNA-bound Sox9-Halo molecules decreases during differentiation from ∼48% in Lgr5+ stem/early progenitor cells to ∼38% in Lgr5- late progenitor/differentiated cells (Fig. 2B; Fig. 7D,E), consistent with Sox9 functioning in ISC-containing crypts^42,51^. In contrast, long-term Sox9 overexpression eventually results in an increased fraction bound of Sox9-Halo molecules (>60%) (Fig. 5C; Fig. 7C-E) underlying a proliferative cell state transition (Fig. 3B,E) and a morphological transition from budding to spherical organoids (Fig. 3A; Fig. 4A). Spheroids lack canonical epithelial cell type markers across lineages (Fig. 4D,E; Fig. S5A) and acquire a gene expression program resembling fetal-like reversion (Fig. 4D,E; Fig. S5B; Fig. 7C-E).

The extension of our previously developed automated SMT pipeline^18^ to the recording of fluorescent cell type markers (Lgr5) allowed us to measure TF diffusion dynamics in a cell type-resolved manner (Fig. 2A), validating our previously established morphology-based broad distinction between ISCs/early progenitors and late progenitors/differentiated cells^18^ (Fig. 1H,I). Parsing Lgr5+/- cells revealed differences in the diffusive behavior of Sox9 correlated with differentiation (Fig. 2; Fig. S2A-D), while a subpopulation of small Lgr5- cells displaying Sox9 diffusion characteristics of Lgr5+ cells (Fig. 2C) potentially constitutes differentiated Paneth cells residing in proliferative centers, testable with additional cell type markers. Similar to our previous study on Hes1^18^ and work in other differentiation contexts^24,68^, Sox9 is characterized by a larger fraction of immobile and hence DNA-bound molecules in the Lgr5+ stem/early progenitor cell subpopulation (Fig. 2B,D; Fig. S2A-D) in which it has been described to function^42,51^. Consistent with Hes1^18^, also functioning during early differentiation, the diffusive behavior of Sox9 is characterized by a rather small average fraction bound of ∼39%, which could be explained by ISCs/early progenitor cells being broadly permissive epigenetic cell states that enable fast transitions to differentiated states^69,70^. In addition, for both Hes1^18^ and Sox9, the overall cell population (Fig. 1E; Fig. S1E) and the proliferative/Lgr5+ population (Fig. 2C; Fig. S2A,C) show a highly heterogenous TF diffusive behavior, suggesting the existence of multiple cell states, consistent with single-cell RNAseq data^71^, and possibly distinct differentiation trajectories for each Lgr5+ cell. Reasons for the latter could include differences in the cellular microenvironment (neighboring cell types), the progression along the same differentiation trajectory (distance from the proliferative center) or different lineages a progenitor cell has committed to. In future work, these could be probed using additional live cell-compatible fluorescent cell type markers in combination with correlative IF for determining end-point cell types and by tracking the same cell during differentiation with time-lapse SMT^18^.

While the differences in Sox9 diffusive behavior were concentration-independent within the probed range of Sox9-Halo transgene expression levels (Fig. S1G; Fig. 2D), long-term overexpression of Sox9 in mSIOs resulted in increased chromatin binding accompanied by a change in organoid morphology towards spherical (Fig. 3A; Fig. 4A). In line with a recent report about Sox9 dosage robustness in the craniofacial context^72^, these observations suggest that the intestinal epithelium can buffer modestly increased Sox9 levels (Fig. 3C) for a limited time to maintain homeostasis (Fig. 4A; Fig. S4D,E). However, long-term Sox9 overexpression appears to break such robustness (Fig. 3A,F,G; Fig. 4A,C,E). While we cannot completely rule out contributions to spheroid formation due to long-term organoid culture, our WT CTRL retaining budding morphology suggests that Sox9 overexpression at least accelerated spheroid formation if it was not the sole cause.

In addition to Sox9 dosage, future research will be required to unravel how other layers of control, such as posttranslational modifications, the availability of tissue-specific co- factors, and chromatin accessibility to gene-regulatory DNA motifs, come into play for Sox9 to achieve its various tissue- and context-dependent functions^34^. In another differentiation system Sox9 has been shown to act both via its capability to access closed chromatin and by competing for epigenetic factors to switch stem cell fates^73^. Related to the Sox9 overexpression-induced spheroids described here, both in the fetal epithelium and upon fetal-like reversion in the adult intestinal epithelium, an increased accessibility was reported for genomic regions enriched for TF binding sites of the SOX and TEAD families^74,75^, both shown to form complexes with Yap/Taz^76^. In the future it will be interesting to employ PAPA-SMT^32,33^ (Fig. 6A,B) to detect the interactions and study the molecular dynamics of such complexes in differentiation and reprogramming contexts. While our PAPA experiments suggest that Sox9 binds to DNA in an oligomerized form (Fig. 6E,G; Fig. S7A), possibly dimers as reported in other contexts^47,48^, this assay cannot determine the degree of Sox9 oligomerization and whether such immobile self-associated Sox9 moieties are all bound at *cis*- regulatory elements of Sox9 target genes. Thus, another possible yet purely speculative scenario could be the formation of gene regulatory Sox9 hubs via weak multivalent interactions mediated by intrinsically disordered regions (IDRs)^77^, testable with IDR mutants of Sox9 in future work.

Constitutive Sox9 expression is a common pathway to cancers through the activation of oncogenic transcription regulators^78,79^, consistent with the gain in proliferative capacity in the Sox9 overexpression-induced spheroids (Fig. 3B,E). However, these spheroids were characterized by a loss in epithelial identity and function towards the activation of a gene expression program resembling that of regenerative fetal-like reversion upon tissue damage^61^ (Fig. 3G; Fig. 4C-E; Fig. S5) consistent with Sox9’s essential role in several developmental pathways^34^. A similar gene expression signature is characteristic of fetal intestinal epithelial cultures in comparison to enteroids generated from adult intestinal cells^80,81^. Hyperproliferative cells and fetal-like gene expression signatures were further found in crypts associated with granulomas resulting from helminth infection^82^ and enriched in the colonic epithelium upon damage induced by dextran sodium sulfate^83^. Similar fetal-like transcriptional programs have been reported upon other injuries to ISCs or the intestinal crypt, such as diphteria toxin-mediated ISC ablation, ionizing radiation, and chemotherapy^80,84–87^. Consistent with the observation of Yap1 activation in Sox9 spheroids (Fig. 4E), such a regenerative response of fetal-like reversion is typically mediated by Yap1 which has a well-established role in driving pattern formation during intestinal regeneration^62–64^. While during an intestinal tissue damage response this transient fetal-like state is exited and intestinal homeostasis is reestablished, it is unclear whether this can be achieved in the Sox9 overexpression-induced spheroid context. To this end, a downregulation of Yap activity together with an upregulation of Wnt and Notch signals to sustain homeostatic ISCs might be required^88^, while retinoid X receptor signaling could also be involved^13^.

The activation of a Yap1-Sox9 circuit is both necessary and sufficient to induce a regenerative fetal-like reversion^61^. However, it has not been reported that Sox9 overexpression alone, whether directly or through intermediate steps, can act itself as a driver of this circuit. Nevertheless, a recent study suggests that Sox9 can promote the nuclear translocation and hence activation of Yap through direct interaction^89^. Further studies will be required to elucidate whether Sox9 overexpression in the intestinal epithelial context directly results in Yap activation and fetal-like reprogramming or whether this involves a more complex cascade of molecular events, including triggering of Yap activation through changes in cell-cell contacts or physical properties of the extracellular matrix^90,91^. In this respect, the long-time scale of several months required for the acquisition of the described fetal-like reprogrammed state will be advantageous for resolving intermediate steps during this organoid morphology transition in multi-pronged, scale-bridging time-course experiments. In addition, it will be interesting to investigate how the epithelial response across scales depends on varying initial Sox9 levels. Furthermore, a reduction in Sox9 expression at various time points during spheroid morphology acquisition will address a potential reversibility of this process. Altogether, while keeping the need of strict control mechanisms to avoid tumor initiation upon constitutive Sox9 expression in mind^78,79^, such studies will inform about the potential of the described Sox9 overexpression-induced fetal-like reversion to be exploited for applications in regenerative medicine.

### Limitations of study

Here, we used Sox9-Halo fusions (Fig. 1B), which were key for live imaging experiments including SMT (Fig. 1C,D). By determining a decreased fraction bound for a Sox9 mutant lacking its DNA binding domain (Fig. 1E-I), we confirmed functionality of Sox9-Halo for DNA binding. Furthermore, we could reproduce a Sox9 overexpression-induced spheroid phenotype with both Halo- and mEGFP-fusions (Fig. 4A). Moreover, we modestly overexpressed transgene-encoded Sox9-Halo in addition to unlabeled endogenous Sox9 (Fig. 3C), necessitating the consideration of the presence of a mixed Sox9 population for data interpretation, which might have resulted in underestimated fractions bound (Fig. 1F,G; Fig. 2B; Fig. 5C; Fig. 7C-E; Fig. S1A,E,G,I; Fig. S2A,B,H,I; Fig. S6A) and oligomerized states (Fig. 6E-H; Fig. S7). Future studies could be improved by endogenous tagging^94^ to investigate Sox9 molecular dynamics at physiological expression levels and to manipulate endogenous levels for interrogating TF dosage effects.

3D mSIOs and 2D EMCs are well characterized differentiation systems recapitulating important features of the intestinal epithelium (Fig. 1A). Nevertheless, they are simplistic *in vitro* models and therefore further studies are required to test the validity of our results in the *in vivo* context and their clinical applicability. SMT deep in tissues remains challenging, and different microscopy techniques capable of imaging single molecules inside thicker specimens^95^ are required for SMT in whole 3D mSIOs. However, our optimization of seeding and imaging conditions enabling HILO-based fast SMT in 3D spheroids (Fig. 7A-C) is an important step towards a broader use in the outer layer of organoid models for connecting multicellular phenotypes with molecular mechanisms.

### Resource availability

#### Lead contact

Further information and requests for resources and reagents should be directed to and will be fulfilled by the lead contact, Nike Walther.

#### Materials availability

Plasmids generated in this study are available upon request with a completed material transfer agreement and with reasonable compensation by the requestor for shipping. There are restrictions to the availability of stable organoid lines generated in this study due to the lack of an external centralized repository for their distribution and our need to maintain the stock.

#### Data and code availability

Microscopy data has been deposited to a public data repository and will be publicly available upon acceptance of this manuscript in a peer-reviewed journal.

RNAseq data has been deposited to the NCBI Gene Expression Omnibus (GEO) under accession number GSE287653 and will be publicly available upon acceptance of this manuscript for publication in a peer-reviewed journal.

Original code has been deposited to GitLab and will be publicly available upon acceptance of this manuscript for publication in a peer-reviewed journal.

Information required to reanalyze the data has been included.

## Acknowledgements

We are grateful to Robert Tjian and Xavier Darzacq for hosting the experimental part of this study in their joint lab at the University of California, Berkeley, and for providing funding (R.T. via the Howard Hughes Medical Institute (34430) and X.D. via a Dynamic Imaging Grant from the Chan Zuckerberg Initiative (Dynamic-0000000091)). We thank current and past members of the Tjian/Darzacq lab for scientific discussions and comments on the manuscript. We are grateful to Qiulin Zhu and Brendan Wu for cloning assistance, Sanchitha Kannabran and Sophia Lim for assistance with plasmid preparations and organoid cultures, Xinyin Lu for plasmid preparations, and Shuang Zheng for assistance with mouse colony maintenance. We thank Thomas Graham for providing code for (PAPA)-SMT automation and analysis. We are thankful to Fred de Sauvage for providing Lgr5::DTR-EGFP mice, Mark Kay for KP1 capsid plasmids, and Luke Lavis for JF dyes. We thank the UC Berkeley Cell Culture Facility supported by The University of California, Berkeley, for providing cell lines. We are grateful to Ophir Klein and current and past members of the Klein lab gut group for sharing protocols and providing a platform for discussion and feedback throughout this study. We would like to thank Fred de Sauvage and Kim Boonekamp for critically reading and commenting on the manuscript. N.W. extends her gratitude to Olaf Stemmann and the Department of Genetics at the University of Bayreuth for hosting her for parts of this study. N.W. acknowledges funding from the Berkeley Stem Cell Center via a Siebel postdoctoral fellowship as well as from the German Research Foundation (DFG) via a Walter Benjamin postdoctoral fellowship (453309976) and a return grant of the Walter Benjamin program (552461306). S.A. was supported by a Donner 160 fellowship. A.C.M. acknowledges support via the California Institute for Regenerative Medicine Training Program (EDUC4-12790).

## Author contributions

N.W. conceptualized, designed, and supervised the study, developed experimental strategies for rAAV-based transgene delivery into 2D EMCs as well as for SMT in 3D spheroids, devised the further development of the automated SMT imaging and analysis pipeline, manually segmented confocal images, analyzed imaging and RNAseq data, and wrote the manuscript (original draft, review/editing). N.W. executed all experiments except for RNA extraction and library preparation for RNAseq, which was performed by C.C., and confocal imaging of one IF condition, which was performed by S.A. S.A. further developed the automated SMT pipeline to accommodate the acquisition of additional fluorescent markers in 2D EMCs and to enable the separate analysis of manually classified SMT data with input from N.W., wrote tailored SMT and image analysis code including statistical analyses, and analyzed a subset of confocal data. C.C. processed RNAseq data and performed initial analyses. S.A. and C.C. discussed data and their interpretation with N.W., provided a subset of raw figure panels as well as methods details, and reviewed and edited the manuscript. G.M.D. designed and cloned DNA constructs. A.C.M provided critical reagents and advice for rAAV-based transgene delivery. N.W. provided funding for this project. All authors agreed on the final version of the manuscript.

## Material and Methods

### Experimental model and subject participant details

#### Mice

C57BL/6J mice (WT) were obtained from The Jackson Laboratory (strain #000664, RRID:IMSR_JAX:000664). Lgr5::DTR-EGFP mice^56^ were kindly provided by Fred de Sauvage (Genentech). WT and Lgr5::DTR-EGFP mice were bred with C57BL/6J mice and housed in an AAALAS-certified level 3 facility on a 14 h light cycle. Pups were weaned 21 d after birth and housed with four littermates per cage. Female offspring was used for mouse small intestinal crypt isolation and organoid generation at an age of 8-16 weeks. All procedures to maintain and use the mice were approved by the Institutional Animal Care and Use Committee of the University of California, Berkeley (IACUC protocol number AUP-2015-09-7988-2).

#### Cell lines

L-Wnt3a cells (CRL-2647, ATCC) were used to produce Wnt3a-conditioned medium. Hek293T cells (CRL-3216, ATCC) were used to generate rAAV vectors for crude rAAV vector preparations, and Hek293T Lenti-X cells (Takara, cat.# 632180) for making lentivirus. Cell lines were obtained via the UC Berkeley Cell Culture Facility. Cell lines and organoids were confirmed to be mycoplasma-free by regular PCR testing.

### Method details

#### Cloning of DNA constructs

For the production of lentivirus to deliver POI-tag transgenes into organoids, the third generation lentiviral pHAGE vector originally developed in the lab of Richard Mulligan^96^ was used together with the second generation lentiviral packaging plasmid psPAX2 (gift from Didier Trono (Addgene plasmid #12260; http://n2t.net/addgene:12260; RRID:Addgene_12260)) and the VSV-G envelope expressing plasmid pMD2.G (gift from Didier Trono (Addgene plasmid #12259; http://n2t.net/addgene:12259; RRID:Addgene_12259)). For constitutive expression of POI-tag under the weak L30 promoter, pHAGE L30 IRES Puro^18^, pHAGE L30 IRES Zeo, and pHAGE L30 IRES Neo backbones were created. The POI-tag transgenes H2B-Halo-V5^18^, V5-Halo-NLS^18^, mSox9-Halo-V5, and mSox9-mEGFP-V5 were cloned into the Puro vector, H2B-mEGFP-V5 was cloned into the Zeo vector, and H2B- mScarletI-V5 was cloned into the Neo vector. For mSox9, the DNA sequence encoding the reference 507aa isoform of UniProt entry Q04887 was cloned (NCBI Gene ID 20682, NM_011448.4). The final pHAGE constructs were confirmed by Sanger sequencing.

For the preparation of crude rAAV to deliver POI-tag encoding transgenes transiently into 2D EMCs, expression cassettes of L30prom_mSox9deltaHMG-NLS-Halo-V5 or L30prom_mSox9-SNAPf-ALFA^97^ were cloned in a pAAV vector derived from pAAV.CMV.Luc.IRES.EGFP.SV40 (gift from James M. Wilson (Addgene plasmid #105533; http://n2t.net/addgene:105533)) to create *cis* plasmids. The helper plasmid pAdDeltaF6 (gift from James M. Wilson (Addgene plasmid #112867; http://n2t.net/addgene:112867)) and the *trans* rep-cap AAV-KP1 plasmid (gift from Mark Kay (Addgene plasmid #206504; http://n2t.net/addgene:206504); ^98^)) were used together with a *cis* plasmid. pAAV bacterial cultures were grown at 30°C and the final rAAV constructs were confirmed by whole plasmid sequencing.

#### Lentivirus preparation

Lentivirus preparation using Hek293T Lenti-X cells (Takara, cat.# 632180) including lentivirus concentration using Lenti-X concentrator (Takara, cat.# 631231) was performed as described^18^.

#### Generation of crude rAAV vector preparations

The generation of crude rAAV vector preparations was based on the protocol described^55^. In brief, 3x10^5^ Hek293T cells (CRL-3216, ATCC) were seeded per well of a 6-well plate (Thermo Fisher Scientific, cat.# 140675) in high-glucose Dulbecco’s modified eagle medium (DMEM) containing GlutaMAX-I (Gibco, cat.# 2537044), supplemented with 10% (vol/vol) fetal bovine serum (FBS; HyClone, cat.# SH30910.03, LOT# AXJ47554), and 1 mM sodium pyruvate (Gibco, cat.# 11360070), and grown to 75-90% confluency. For transfection of a 6-well, a plasmid mixture of 1.3 µg *trans* rep-cap AAV-KP1 plasmid, 2.6 µg helper plasmid pAdDeltaF6, and 1.3 µg gene of interest (GOI)-encoding *cis* plasmid (see “Cloning of DNA constructs”) was prepared in serum-free DMEM to 100 µL per well. After addition of 5.2 µL 1 µg/µL polyethylenimine (PEI) hydrochloride MAX (Polysciences, cat. # 24765-1) in distilled H_2_O (pH 7.1, filter sterilized), the mixture was pulsed 10-15x on a vortexer. Following incubation for 15 min at RT, 1.9 mL serum-free DMEM was added and mixing was performed by gently pipetting up and down two times. After removal of the culture medium, 2 mL of transfection mixture was added per well and the plate was incubated at 37°C and 5% CO_2_ for 72 h. The 6-well plate was frozen at -80°C for 30 min, followed by thawing at 37°C for 30 min. Freezing and thawing was repeated for a total of three cycles. Each well was mixed by pipetting and the cell lysate was transferred to a 2 mL tube. To remove cell debris, centrifugation was performed at 15,000 x *g* for 15 min at RT. The supernatant (SN) was removed and transferred to a new 2 mL tube. This crude rAAV vector preparation was stored at 4°C for several weeks before being used for transduction of 2D EMCs.

#### Mouse small intestinal crypt preparation for growing small intestinal organoids

Mouse small intestinal crypt preparations for growing small intestinal organoids were performed as described^18^ with the exception that female mice at an age of 8-16 weeks were used for this study.

#### Mouse small intestinal organoid culture

Mouse small intestinal organoids were grown in Matrigel domes (90% (vol/vol) Matrigel Matrix for Organoid Culture (Corning, cat.# 08774406)) and cultured in IntestiCult Organoid Growth Medium (Mouse) (Stem Cell Technologies, cat.# 06005) supplemented with 1% (vol/vol) Penicillin/Streptomycin (Pen/Strep; Gibco, cat.# 15070063) as described^18^. mSIOs were passaged every week using Gentle Cell Dissociation Reagent (Stem Cell Technologies, cat.# 100-0485) with one to two media changes in between depending on mSIO density except for Sox9 spheroids which required at least two media changes due to their increased growth/proliferation rate.

#### Preparation of Wnt3a-conditioned medium

Wnt3a-conditioned medium was prepared as described before^18^ using L-Wnt3a cells (CRL-2647, ATCC) and Advanced DMEM/F-12 (ADMEM; Gibco, cat.# 12634028) supplemented with 10% (vol/vol) FBS, 1% (vol/vol) GlutaMAX (Gibco, cat.# 35050061), 10 mM HEPES (Sigma-Aldrich, cat.# H0887-20mL), 1 mM N-acetyl-L-cysteine (Sigma-Aldrich, cat.# A9165-5G; 500 mM stock in ddH_2_O, sterile-filtered).

#### Lentiviral transduction of mouse small intestinal organoids and selection of stable organoid lines

Lentiviral transduction of mSIOs was performed as described before^18^. In brief, mSIOs were seeded in pre-transduction medium consisting of 50% (vol/vol) IntestiCult without Pen/Strep and 50% (vol/vol) Wnt3a-conditioned medium, supplemented with 100 mM nicotinamide (Sigma, cat.# N0636-100G; 1M stock in ddH_2_O), 10 μM ROCK inhibitor Y-27632 (Stem Cell Technologies, cat.# 72304; 10 mM stock in ddH_2_O), and 2.5 μM CHIR99021 (Stem Cell Technologies, cat.# 72052; 10 mM stock in dimethylsulfoxide (DMSO; Sigma-Aldrich, cat.# D2650)). 4-5 d post-seeding, mSIOs were broken down into single cells using TrypLE Express (Thermo Scientific, cat.# 12604013) prior to spinoculation (1 h at 37°C and 600 g) with lentivirus in pre-transduction medium containing 10 μg/mL polybrene Millipore Sigma, cat.# TR-1003-G) and incubation for 6 h at 37°C. Transduced cells were seeded in 90% (vol/vol) Matrigel Matrix for Organoid Culture in IntestiCult Organoid Growth Medium supplemented with 1% (vol/vol) Pen/Strep and grown in pre-transduction medium. 2-3 d post-transduction, the medium was exchanged to pre-transduction medium containing selection antibiotics (2 μg/mL puromycin (Thermo Scientific, cat.# A11138-03)) and the selection medium was exchanged every 2-3 d until large, selected spheroids were obtained. Upon passaging, selected spheroids were cultured in IntestiCult supplemented with 10 μM ROCK inhibitor Y-27632 and 2.5 μM CHIR99021 including selection antibiotics. After 2-3 d the ROCK inhibitor was removed, followed by the removal of CHIR99021 after another 2-3 d to culture selected stable organoid lines as described before including selection antibiotics.

Established stable mSIO lines were cryopreserved in CryoStor CS10 (Stem Cell Technologies, cat.# 07931) as described before^18^ and stored in liquid nitrogen.

Except for Sox9 spheroids and when specifically noted otherwise, mSIOs were used for experiments until 8 weeks after line establishment or an organoid aliquot cryopreserved at an early passage was thawn as described before^18^.

> For stable Sox9-Halo or Sox9-mEGFP organoids (WT background), a second transduction was performed at P15(P12) according to the same protocol to achieve double stable organoid lines co-expressing H2B-mEGFP or H2B-mScarletI, whereby antibiotic selection was performed using 40 µg/mL Zeocin (Fisher Scientific, cat.# NC9002627) or 200 µg/mL Geneticin (Thermo Fisher Scientific, cat.# 10131027; neomycin resistance), respectively.

#### Brightfield imaging of organoid lines and 2D enteroid monolayer cultures

Brightfield (BF) imaging of mSIOs or 2D EMCs growing on 24-well culture plates (Corning, cat.# 353047) or 8-well Labteks II #1.5 (Nunc, cat.# 12-565-338) to record 3D/2D morphology was performed on an EVOS M5000 Imaging System (Invitrogen) microscope using 10x (Fig. 3A) or 4x (Fig. 4A) air objectives 5 d post-seeding.

#### Confocal imaging of Sox9-Halo organoid lines

For confocal imaging of Sox9-Halo organoid lines, POI-Halo organoids were seeded in 10-20 μL 90% Matrigel in IntestiCult droplets into 8-well Labteks II #1.5 (Nunc, cat.# 12-565-338) and overlaid with 500 μL IntestiCult. After 3 d the medium was exchanged. 5 d post-seeding, organoids were stained with 500 nM HTL-JF635 (kind gift from Luke Lavis) in DMEM/F-12 without phenol red (Gibco, cat.# 11039-021) for at least 1 h and imaged 1-6 h after staining.

Confocal imaging of 3D mSIOs was performed on a Yokogawa CSU-W1 SoRa spinning disk with a Nikon Ti2 inverted microscope operated by the NIS Elements AR 5.42.03 software (Nikon). The microscope was equipped with a temperature- and CO_2_- controlled incubation chamber (Okolab), and the temperature was set to 37°C and the CO_2_ to 5% for imaging of live organoids. For combined BF/Sox9-Halo organoid images (Fig. 1C, Fig. S3A), acquisition was performed using an Apo LWD 40× WI λS DIC N2 water immersion objective (N.A. 1.15; Nikon). For imaging Sox9-Halo-JF635 and Lgr5- DTR-GFP (budding; Fig. 1C) or H2B-mEGFP (spheroid; Fig. S3A), JF635 was excited with 640 nm (laser at 80% (budding) or 40% (spheroid)) and detected with Hamamatsu Orca Flash 4.0 cameras at 200 ms exposure using a 561 nm long-pass (LP) filter. Consecutively, the widefield fluorescence imaging modality was used for BF detection with a Hamamatsu Orca Flash 4.0 camera at 100 ms exposure. For each modality, multiple *z* planes (*xy* pixel size: 0.1625 × 0.1625 μm; *z* interval 0.4 μm, number of *z* slices varying depending on the thickness of organoids) were imaged using a NIDAQ *z* piezo. One selected *z* plane is shown in Fig. 1C and Fig. S3A.

#### Generation of 2D enteroid monolayer cultures from 3D mouse small intestinal organoids and seeding on imaging dishes

2D EMCs were derived from 3D mSIOs as described^10,18^. In brief, a 1:40 (vol:vol) mixture of Matrigel Growth Factor Reduced Basement Membrane Matrix (Corning, cat.# 356231) in DMEM/F-12 without phenol red was used for coating imaging dishes by incubation for at least 30 min up to one week in the cell culture incubator prior to seeding. 100-500 μL of 1x10^6^ cells/mL mSIO-derived single-cell suspensions were seeded in plating medium (IntestiCult containing 20 μM ROCK inhibitor and 3 μM CHIR) to achieve a total of 500 μL/well of an 8-well Labtek II #1.5. 1-2 d after seeding and every other day thereafter the medium was exchanged to IntestiCult.

#### Confocal imaging of POI-Halo enteroid monolayer cultures

2D EMCs were seeded and cultured as described above. 5 d post-seeding, EMCs were stained with 200 nM HTL-JFX549 (kind gift from Luke Lavis) in DMEM/F-12 without phenol red for 30 min. After two washes with DMEM/F-12 without phenol red for 15 min each, the medium was exchanged one more time to DMEM/F-12 without phenol red before proceeding with imaging.

Confocal imaging of live 2D EMCs (Fig. 1C, Fig. S3A) was performed on an LSM900 Airyscan 2 laser-scanning microscope with an inverted Axio Observer.Z1 / 7 operated by the ZEN 3.1 blue software (ZEISS). The microscope was equipped with a temperature- and CO_2_-controlled incubation chamber (Zeiss/PeCon), and the temperature was set to 37°C and the CO_2_ to 5%. Images were acquired using a Plan-Apochromat 40×/N.A. 1.3 Oil DIC (UV) VIS-IR M27 oil-immersion objective (ZEISS). One *z* plane was imaged consecutively in the 561 nm and 488 nm channels (*xy* pixel size: 0.092 × 0.092 μm; 1.21 μs pixel dwell time; bidirectional scanning; 4-times averaging). JFX549 was excited with 561 nm (diode (SH) laser at 2.0%) and detected with GaAsP (spectral gallium arsenide) detectors at 566-635 nm. Green fluorescence was excited with 488 nm (diode laser at 4.5% for Lgr5-DTR-GFP (Fig. 1C) or 0.1% for H2B-mEGFP (Fig. S3A)) and detected with a multialkali-photomultiplier (MA-PMT) detector at 410-545 nm.

Images acquired as described above were also used for Sox9-Halo transgene expression level quantification in Sox9 budding organoids *versus* spheroids (Fig. 3C, right).

For confocal imaging of rAAV-transduced conditions, 2D EMCs were prepared and stained as described under “Transient transduction of 2D enteroid monolayer cultures using crude rAAV vector preparations for SMT and PAPA-SMT” and imaged on a spinning disk confocal microscope described under “Confocal imaging of Sox9-Halo organoid lines” using an Apo LWD 40× WI λS DIC N2 water immersion objective (N.A. 1.15; Nikon). For imaging Sox9ΔHMG-Halo-JFX650 (Fig. S1D), JFX650 was excited with 640 nm (laser at 40%) and detected with a Hamamatsu Orca Flash 4.0 camera at 500 ms exposure using a 640 nm LP filter. For imaging Sox9-Halo-JFX549/Sox9- STL-JFX650 (Fig. 6C) and Sox9-Halo-JFX549/JFX650-STL-SNAPf-3xNLS (Fig. 6D), JFX650 was excited with 640 nm (laser at 50%) and JFX549 was excited with 561 nm (laser at 50%). Fluorescence was detected with Hamamatsu Orca Flash 4.0 cameras at 300 ms exposure using a 640 nm LP filter. Multiple *z* planes (*xy* pixel size: 0.1625 × 0.1625 μm; 21 *z* slices at *z* interval 0.4 μm) were imaged using a NIDAQ *z* piezo. One selected *z* plane is shown in Fig. S1D and Fig. 6C,D.

For confocal imaging of cell division phenotypes in 2D EMCs derived from Sox9-Halo spheroids co-expressing H2B-mEGFP (Fig. S3B), EMCs were prepared as described above. Instead of Sox9-Halo, staining for microtubules was performed by adding 500 nM SiR-tubulin and 10 µM Verapamil (SiR-tubulin kit; Cytoskeleton, cat.# CY-SC002) in DMEM/F-12. After incubation at 37°C for 1 h, live imaging was performed on a spinning disk confocal microscope described under “Confocal imaging of Sox9-Halo organoid lines” using an Apo LWD 40× WI λS DIC N2 water immersion objective (N.A. 1.15; Nikon). SiR-tubulin was excited with 640 nm (laser at 50%) and H2B-mEGFP was excited with 488 nm (laser at 5%). Fluorescence was detected with Hamamatsu Orca Flash 4.0 cameras at 200 ms exposure using a 561 nm LP filter. Multiple *z* planes (*xy* pixel size: 0.1083 × 0.1083 μm; 37 *z* slices at *z* interval 0.3 μm) were imaged using a NIDAQ *z* piezo. One selected *z* plane is shown in Fig. S3B.

#### IF of 2D enteroid monolayer cultures and confocal imaging

IF of 2D EMCs and confocal imaging was performed 5 d post-seeding as described^18^. In brief, EMCs were fixed with 4% paraformaldehyde (PFA; Electron Microscopy Sciences, cat.# EMS14710) in PBS for 20 min at RT, washed three times with PBS for 5 min and stored at 4°C or directly subjected to IF staining. Permeabilization was performed with 0.5% Triton X-100 (TX-100; Sigma-Aldrich, cat.# T9284) in PBS for 1 h. Following three washes for 5 min each with PBS, blocking was performed in 3% (vol/vol) donkey serum (Sigma-Aldrich, cat.# D9663) in 0.1% (vol/vol) TX-100 in PBS (blocking buffer) for 4 h at RT or ON at 4°C. Incubation with the primary antibody (AB) in blocking buffer was performed ON at 4°C in a humidified chamber. Following three washes in blocking buffer for 5 min each, incubation with the secondary AB was performed for 1 h at RT. Following three washes with PBS for 5 min each, DNA was stained with 1 μg/mL Hoechst 33342 (Thermo Scientific, cat.# H3570) in PBS for 10- 15 min, followed by another three washes with PBS for 5 min each. Immunostained samples were either imaged directly or sealed with parafilm and stored for short-term at 4°C prior to imaging.

The following primary ABs and dilutions were used: rabbit anti-Sox9 (Abcam, cat.# ab185966; 1:200), rabbit anti-Ki67 (Abcam, cat.# ab16667; 1:200), rabbit anti-Olfm4 (Cell Signaling, cat.# 39141; 1:50), rabbit anti-Lysozyme (Agilent, cat.# A009902-2; 1:300), sheep anti-Dll1 (Biotechne, cat.# AF3970; 1:20), rabbit anti-Aldolase B/C (Abcam, cat.# ab75751; 1:50), rabbit anti-Sca-I (Abcam ab124688, cat.# ab124688; 1:50), rabbit anti-Yap (Cell Signaling, cat.# 14074S; 1:100), rabbit anti-ß-catenin (Abcam, cat.# ab32572; 1:250), rabbit anti-E-Cadherin (Abcam, cat.# ab40772; 1:1000), rabbit anti-EpCAM (Thermo Fisher Scientific, cat.# MA5-35283; 1:100). A secondary donkey anti-rabbit-AlexaFluor568 AB (Thermo Fisher Scientific, cat.# A10042) was used. For F-actin staining, incubation with a 1x phalloidin-AlexaFluor568 (Thermo Fisher Scientific, cat.# A12380) staining solution in PBS was performed for 1 h at RT instead of incubation with primary/secondary ABs.

Confocal imaging of immunostained monolayers (Fig. 3B; Fig. 4E; Fig. S3C) was performed on an LSM900 Airyscan 2 laser-scanning microscope with an inverted Axio Observer.Z1 / 7 operated by the ZEN 3.1 blue software (ZEISS) at RT. Images were acquired using a Plan-Apochromat 40×/ N.A. 1.3 Oil DIC (UV) VIS-IR M27 oil-immersion objective (ZEISS). One *z* plane was imaged in the 561 nm and 405 nm channels (*xy* pixel size: 0.11 × 0.11 μm; 2.06 μs pixel dwell time; bidirectional scanning; 4-times averaging). AlexaFluor568 was excited with 561 nm (diode (SH) laser at 0.5- 5.0% depending on the brightness of the immunostaining) and detected with MA-PMT detector at 576-700 nm. Hoechst was excited with 405 nm (diode laser at 0.2% except for Aldolase condition (0.5%)) and detected with MA-PMT detector at 410-493 nm. Laser excitation powers were kept constant across all samples per immunostaining condition.

Images immunostained for Sox9 and Ki67 and acquired as described above were also used for quantifications of the total Sox9 expression level (Fig. 3C left) or the fraction of proliferative cells (Fig. 3E) in Sox9 spheroids *versus* WT organoids.

#### Image analysis of Sox9 immunostained 2D enteroid monolayer cultures to estimate total Sox9 expression levels

Hoechst-stained nuclei in confocal images of WT or Sox9_spheroid 2D EMCs immunostained for Sox9 (see “IF of 2D enteroid monolayer cultures and confocal imaging”) were manually segmented using ImageJ^99^. Following conversion of manual segmentation masks into Cellpose^101^ compatible masks, mean Sox9 intensities were extracted using the Scikit-image library^100^ and compared between WT and Sox9_spheroid conditions (Fig. 3C left). For the WT condition, cells were classified into Sox9-positive and Sox9-negative and analyzed separately to distinguish on-target from background fluorescence. Quantifications were based on 3 images with 456 cells (162 Sox9+, 294 Sox9-) for WT and 3 images with 159 cells for Sox9_spheroid from one IF experiment, whereby cells in each image were aggregated and the mean was calculated.

#### Image analysis of HaloTag-stained live Sox9-Halo 2D enteroid monolayer cultures to estimate Sox9-Halo transgene expression levels

Nuclei in confocal images of WT or Sox9_spheroid live 2D EMCs (see “Confocal imaging of POI-Halo enteroid monolayer cultures”) were manually segmented using ImageJ^99^ based on the predominant nuclear localization of HTL-JFX549-labeled Sox9- Halo. Following conversion of manual segmentation masks into Cellpose^101^ compatible masks, mean Sox9-Halo intensities were extracted using the Scikit-image library^100^ and compared between Sox9_budding and Sox9_spheroid conditions (Fig. 3C right). Quantifications were based on 3 images with 864 cells for Sox9_budding and 3 images with 537 cells for Sox9_spheroid from one live-cell imaging experiment, whereby cells in each image were aggregated and the mean was calculated.

#### Image analysis of Ki67-immunostained 2D enteroid monolayer cultures to quantify the fraction of proliferative cells

Hoechst-stained nuclei in confocal images of 2D EMCs immunostained for Ki67 (see “IF of 2D enteroid monolayer cultures and confocal imaging”) were manually segmented using ImageJ^99^ and classified into Ki67-positive and Ki67-negative. Following conversion of manual segmentation masks into Cellpose^101^ compatible masks, mean Ki67 intensities were extracted using the Scikit-image library^100^ and compared between WT and Sox9_spheroid conditions (Fig. 3E). Quantifications were based on 7 images with 752 cells for WT and 7 images with 648 cells for Sox9_spheroid from one IF experiment, whereby cells in each image were aggregated and the mean was calculated.

#### Image analysis of enteroid monolayer cultures to quantify cellular morphological features

To quantify the cellular morphological features nuclear area and nearest-nuclei- distance (Fig. 3D), WT and Sox9_spheroid confocal images of live or immunostained 2D EMCs were pooled, including the three experiments above (determination of total Sox9 expression level (IF), Sox9-Halo transgene expression level (live), and proliferation frequency (IF)). Based on the manually segmented and converted nuclear masks, the Scikit-image library^100^ was used to compute the nuclear area and the mean distance from the centroid of a nucleus to the two nearest centroids of its neighboring nuclei (nearest-nuclei-distance). Quantifications were based on 23 images with 3029 cells for WT and 30 images with 3100 cells for Sox9_spheroid from a total of 4 experiments, whereby cells in each image were aggregated and the mean was calculated. Multinucleated cells were excluded from this analysis.

#### Classification and quantification of cell division errors in 2D enteroid monolayer cultures

To classify and quantify aberrant cell division phenotypes in WT, Sox9_budding, and Sox9_spheroid conditions, confocal images of live or immunostained 2D EMCs were used. The cell division phenotypes (a) two nuclei fused, (b) three nuclei fused, (c) micronuclei, and (d) multiple fragmented nuclei or micronuclei were classified according to the examples in Fig. S3B (left) and their occurrence per total number of cells per image was quantified. Quantifications were based on 13 images with 1821 cells for WT, 8 images with 1626 cells for Sox9_budding, and 17 images with 1756 cells for Sox9_spheroid from a total of 4 experiments, whereby cells in each image were aggregated and the mean was calculated. Not all conditions were included in each experiment.

#### Preparation of 2D enteroid monolayer cultures for SMT and PAPA-SMT

2D EMCs derived from stable Sox9-Halo mSIO lines were seeded and cultured as described above. 2-6 d post-seeding (depending on the degree of monolayer confluency and the formation of crypt- and villus-like morphological characteristics), monolayers were stained for SMT with 50 nM HTL-JFX549 (bulk labeling for nuclear segmentation and feature extraction; kind gift from Luke Lavis) and 1 nM HTL-JFX650 (sparse labeling for SMT; kind gift from Luke Lavis) in DMEM/F-12 without phenol red for 15 min. Following two washes with DMEM/F-12 without phenol red for 15 min each, medium was replaced one more time to DMEM/F-12 without phenol red before proceeding with imaging.

#### Transient transduction of 2D enteroid monolayer cultures using crude rAAV vector preparations for SMT and PAPA-SMT

2D EMCs derived from WT mSIOs (for SMT of Sox9ΔHMG-Halo) or stable Sox9-Halo budding organoid or spheroid lines (for PAPA-SMT) were seeded and cultured as described above. 1-3 d post-seeding, the medium was replaced with a transduction mixture of 250 µL crude rAAV preparation and 250 µL plating medium. On the next day, the transduction mixture was removed. As best transgene expression was reached about 2 d post-transduction, IntestiCult w/o Pen/Strep and selection antibiotics was added for incubation for another day before staining for SMT with 50 nM HTL-JFX549 and 1 nM HTL-JFX650 in DMEM/F12 without phenol read for 15 min at 37°C. For PAPA-SMT, staining with 50 nM HTL-JFX549 together with 50 nM STL- JFX650 (Sox9-SNAPf) or 5 nM STL-JFX650 (SNAPf-3xNLS) in IntestiCult w/o Pen/Strep and selection antibiotics was performed ON at 37°C before PAPA-SMT on the next day. Upon labeling for SMT or PAPA-SMT, two washes for 30 min each in DMEM/F12 without phenol red were performed prior to imaging in DMEM/F12 without phenol red.

#### Preparation of Sox9-Halo spheroids for SMT

For SMT of 3D spheroids, Sox9-Halo spheroids were seeded in 10-20µL droplets in 50% Matrigel in IntestiCult in 8-well Labteks II #1.5 and cultured as described before. Spheroids were grown until they were almost or even slightly touching the glass bottom of the culture dish, resulting in a restricted area within spheroids in which a layer of cells was oriented almost in parallel to the glass bottom. Such a configuration, required for successful HILO-based SMT, was typically reached 5 d post-seeding. Staining was performed with 50 nM HTL-JFX549 and 1-2.5 nM HTL-JFX650 in DMEM/F12 without phenol red for 1 h at 37°C. Following four washes in DMEM/F12 without phenol red for 30 min each and replacing the medium one more time with fresh DMEM/F12, SMT was performed.

#### TIRF microscope for HILO-based fast SMT and PAPA-SMT

All SMT and PAPA-SMT experiments were performed using a custom-built microscope as previously described^26^. In brief, a Nikon Ti microscope was equipped with a 100×/N.A. 1.49 oil-immersion TIRF objective (Nikon apochromat CFI Apo TIRF 100× Oil), a motorized mirror, a perfect focus system, an EM-CCD camera (Andor iXon Ultra 897), a laser launch with 405 nm (140 mW, OBIS, Coherent), 488 nm, 561 nm and 639 nm (1 W, Genesis Coherent) laser lines, and an incubation chamber maintaining a humidified atmosphere with 5% CO_2_ at 37°C. The NIS-Elements software (Nikon) was used for controlling all microscope, camera and hardware components. Imaging was performed at laser power densities of approximately 52 W/cm^2^ for 405 nm (violet), 91 W/cm^2^ for 488 nm (blue), 100 W/cm^2^ for 561 nm (green), and 2.3 kW/cm^2^ for 639 nm (red).

#### Microscope automation for SMT and PAPA-SMT

A custom-built inverted Nikon Ti microscope described above was automated using code written in Python and the NIS Elements Macro Language as previously described^18,29,33^. In brief, the microscope stage rastered in a grid-like pattern to move the sample. At each grid position, a 512 px x 512 px (6710.9 µm^2^) image was recorded in the densely labeled JFX549 channel or in the H2B-mEGFP channel for Sox9- Halo_spheroid. Segmentation of nuclei was performed using the pre-trained Versatile Fluorescent Nuclei model of the Python package StarDist^102^. After randomly choosing one of the nuclei within a range of user-defined brightness and size parameters, the stage was moved to center it in the large FOV. The FOV was then resized to contain a smaller 150 px x 150 px (576 µm^2^) square (small FOV; zoom-in) centered on the chosen nucleus. Images of the large and small FOVs were recorded in multiple channels corresponding to the fluorophores in the sample (GFP, JFX549, JFX650). JFX650 fluorophores in the small FOV were pre-bleached with red light at 639 nm to achieve sparse and thus trackable single molecules, following which an illumination sequence for fast SMT or PAPA-SMT was executed. On-the-fly localization density assessment was performed using quot (https://github.com/alecheckert/quot; ^103^). Microscope automation was based on our previously published code (^18^; https://gitlab.com/tjian-darzacq-lab/walther_2dmsios_automatedsmt_2024/-/tree/main/Microscope_Automation_Scripts?ref_type=heads).

#### Fast SMT of POI-Halo in enteroid monolayer cultures

For imaging in GFP, JFX549 or JFX650 channels, laser powers were set to 100 mW for 488 nm, 110 mW for 561 nm, and to 1100 mW for 639 nm. The exposure time was set between 20 ms and 200 ms with a laser excitation power between 1.35% and 20% to achieve segmentable bulk labeling of POI-Halo depending on its expression level and to avoid saturation. Imaging conditions were kept constant across experiments for the same POI-Halo condition. Semrock 536/40 nm, 593/40 nm or 676/37 nm bandpass filters, respectively, were used.

For fast SMT experiments, the following bleaching durations in the 639 nm channel (100% laser excitation power) prior to executing an illumination sequence were used: H2B – 15s, NLS – 10s, Sox9_budding – 15s or 10s, Sox9_spheroid – 15s, Sox9ΔHMG – 15s.

The triggered illumination sequence consisted of the following phases at a frame rate of 7.48 ms/frame:

1. Imaging: 5000 frames of red light (639 nm, one 2 ms stroboscopic pulse per frame) with 5% 405 nm reactivation (pulsed during the 0.48 ms camera transition time between 7 ms detection windows)

2. Imaging: 5000 frames of red light (639 nm, one 2 ms stroboscopic pulse per frame) with 10% 405 nm reactivation (pulsed during the 0.48 ms camera transition time between 7 ms detection windows)

To reduce the motion blur of moving molecules^104^, red illumination was restricted to stroboscopic pulses during each frame of a single-molecule movie.

#### Fast SMT of Sox9-Halo in Sox9-Halo spheroids

Fast SMT in 3D spheroids was performed in analogy to SMT in 2D EMCs with the exception that cells within spheroids that were close enough (10-20 µm) to the glass bottom of the cover slip to be reachable by HILO illumination were manually selected and centered within the FOV before triggering a sequence of epifluorescence imaging and SMT.

The following modified SMT illumination sequence was used:

1. As the bleaching efficiency in the 639 nm channel depended on the distance of the cell from the glass bottom of the culture dish, its duration had to be adjusted manually for each cell and was carried out until a trackable density of single-molecule localizations was achieved.
2. Embedding in Matrigel and the mostly larger distance of selected cells in 3D spheroids from the glass bottom of the culture dish in comparison to 2D EMCs required a higher 405 nm reactivation. The triggered illumination sequence thus consisted of the following phases at a frame rate of 7.48 ms/frame:

1. Imaging: 5000 frames of red light (639 nm, one 2 ms stroboscopic pulse per frame) with 0% 405 nm reactivation (primarily used to estimate background reactivation and non-specific fluorescence to judge suitability of selected cell for SMT)
2. Imaging: 5000 frames of red light (639 nm, one 2 ms stroboscopic pulse per frame) with 20% 405 nm reactivation (pulsed during the 0.48 ms camera transition time between 7 ms detection windows)
3. Imaging: 5000 frames of red light (639 nm, one 2 ms stroboscopic pulse per frame) with 100% 405 nm reactivation (pulsed during the 0.48 ms camera transition time between 7 ms detection windows)

#### PAPA-SMT of POI-Halo in enteroid monolayer cultures

Imaging for PAPA-SMT was performed as described for fast SMT.

For all PAPA-SMT conditions, an initial bleaching in the 639 nm channel (100% laser excitation power) was performed for 15s prior to executing an illumination sequence.

For PAPA-SMT experiments, the illumination sequence consisted of 5 cycles of the following phases at a frame rate of 7.48 ms/frame:

1. Imaging: 250 frames of red light (639 nm), one 2 ms stroboscopic pulse per frame, recorded
2. DR pulse: 10 frames of violet light (405 nm), continuously during 7 ms detection window, recorded
3. DR imaging: 250 frames of red light, one 2 ms stroboscopic pulse per frame, recorded
4. Bleaching: 200 frames of red light, continuously during 7 ms detection window, not recorded
5. Imaging: 250 frames of red light, one 2 ms stroboscopic pulse per frame, recorded
6. PAPA pulse: 100 frames of green light (561 nm), continuously during 7 ms detection window, recorded
7. PAPA imaging: 250 frames of red light, one 2 ms stroboscopic pulse per frame, recorded
8. Bleaching: 200 frames of red light (639 nm), continuously during 7 ms detection window, not recorded

#### SMT and PAPA-SMT data processing and analysis

##### Image analysis for SMT and PAPA-SMT

Image analysis for SMT and PAPA-SMT was performed as previously described^18^. In brief, nuclei in epifluorescence images corresponding to SMT or PAPA-SMT movies were segmented using StarDist^102^. Segmented nuclei were subjected to manual QC for downstream analysis using CellPicker (^18^; https://github.com/tgwgraham/basic_PAPASMT_analysis). Hereby, nuclei were filtered based on their intensities in each channel and a QC was performed to exclude nuclei with erroneous segmentation masks, unusual textures, or masks corresponding to segmented autofluorescent debris. In addition, CellPicker enabled the manual classification of cells based on the expression of the additionally recorded Lgr5-DTR-GFP marker for stem/early progenitor cells. For each category, the cell morphological parameter nuclear area as well as the mean POI intensity in different channels as a proxy for relative POI expression level were computed for each of the chosen nuclei using the Python library Scikit-image^100^ and the nuclear masks generated by StarDist. For nuclei in FOVs with three or more nuclei, the distance from the centroid of the nucleus of interest to the centroid of every other nucleus in the FOV was computed and the mean distance of the centroid of the nucleus of interest to its two nearest neighbors was recorded (nearest-nuclei-distance).

##### SMT data processing and analysis

SMT movies were processed using quot (https://github.com/alecheckert/quot; ^103^), which identifies single-molecule localizations and generates trajectories, as previously described^18^. The following settings were used: [filter] start = 0; method = ‘identity’; chunk_size = 100; [detect] method = ‘llr’; k=1.0; w=9, t=18; [localize] method = ‘ls_int_gaussian’, window size = 9; sigma = 1.0; ridge = 0.001; max_iter = 10; damp = 0.3; camera_gain = 109.0; camera_bg = 470.0; [track] method = ‘conservativeeuclidean’; pixel_size_µm = 0.160; frame interval = 0.00748; search radius = 1; max_blinks = 0; min_IO = 0; scale – 7.0. To construct trajectories, a conservative method was used, in which only trajectories with unambiguously assigned localizations were considered. Trajectories were assigned to cells using the StarDist-generated nuclear masks. Diffusion coefficient distributions were obtained using the state array method of saSPT (https://github.com/alecheckert/saspt; ^103^). Here, a regular Brownian motion with a normally distributed, mean-zero localization error (RBME) model was fit to populations of single-molecule trajectories in each cell to obtain a state array, which consists of posterior occupations of states defined by their localization errors and diffusion coefficients. To obtain diffusion spectra, these state array distributions were marginalized on the diffusion coefficients. Mean diffusion spectra for each POI were constructed by averaging over single-cell diffusion spectra, whereby cells were weighted by the number of trajectories. The fraction of bound trajectories for a POI/condition was defined as the fraction of trajectories with diffusion coefficients below 0.15 µm^2^/s (fraction bound). Fractions of bound trajectories were also computed for individual nuclei based on the single-cell diffusion spectra to correlate them with extracted morphological features, such as nuclear area and nearest-nuclei-distance, as well as the POI expression level. To perform these processing and analysis steps, we used our previously published^18^ custom-written Jupyter notebook for an all-in-one cell level-based diffusion analysis and correlation with extracted cellular features (https://gitlab.com/tjian-darzacq-lab/walther_2dmsios_automatedsmt_2024/-/tree/main/SMT_analysis_scripts?ref_type=heads).

For fast SMT data in Sox9-Halo spheroids, a filter was set to include only cells with at least 100 non-singlet trajectories for downstream analysis.

##### Cluster analysis based on single-cell diffusion spectra

Cluster-based analyses of single-cell diffusion spectra derived from fast SMT data were performed as previously described^18^: The Jensen-Shannon distance as the distance metric^105^ was used to compute a matrix of pairwise distances between pairs of single-cell diffusion spectra. Cells with complete linkage were then hierarchically clustered using the AgglomerativeClustering class of the Scikit-learn Python library^106^. The over- and under-representation of cells from different conditions in different clusters was quantified with a *p*-value computed from the hypergeometric distribution. To account for multiple comparisons, the significance threshold was Bonferroni-corrected. The option for diffusion-based cluster analysis is included in our previously published^18^ custom-written Jupyter notebook (https://gitlab.com/tjian-darzacq-lab/walther_2dmsios_automatedsmt_2024/-/tree/main/SMT_analysis_scripts?ref_type=heads).

##### PAPA data processing and analysis

PAPA movies were processed and analyzed as described for SMT with the following specifications: All trajectory segments occurring within the first 30 frames after pulses of 561 nm light (PAPA trajectories) or 405 nm light (DR trajectories) were extracted using custom MATLAB code reported previously (^18,32^; https://github.com/tgwgraham/basic_PAPASMT_analysis). PAPA and DR trajectories were then separately analyzed similar to SMT data.

#### Bulk RNAseq

##### Organoid collection for RNAseq

For bulk RNAseq experiments in biological triplicates, mSIOs were used at the passages indicated in Fig. 4A plus/minus two passages. Organoids were seeded into 6 wells of a 24-well plate per line and cultured as described before. 5 d post-seeding, the medium was removed and organoids from the 6 wells were harvested with 1 mL TRIzol (Thermo Fisher Scientific, cat.# 15596026) into low-binding 1.7 mL tubes (Sorenson, cat.# 39640T) by rigorously pipetting up and down with a 1 mL pipette to dislodge and homogenize the organoid-containing Matrigel domes. Organoid-TRIzol samples were directly stored at -20°C until RNA extraction.

##### RNA extraction and poly-A RNAseq library preparation

Poly-A RNAseq was performed in three biological replicates per condition. Total RNA was extracted with TRIzol according to the manufacturer’s instructions by performing the optional centrifugation step of the lysates (5 min at 12,000 g at 4–10°C) and an additional wash with one volume of chloroform after the recommended phenol:chloroform extraction (UltraPure Phenol:Chloroform:Isoamyl Alcohol, 25:24:1, v/v, cat.# 15593-01). RNA was quantified by spectrophotometer (NanoDrop, ThermoFisher Scientific) and checked for integrity by capillary electrophoresis (Fragment Analyzer, Agilent). 100-500 ng of total RNA were subjected to poly-A purification and library preparation with the NEBNext Poly(A) mRNA Magnetic Isolation Module (NEB, cat.# E7490S) in combination with the NEBNext Ultra II RNA Library Prep Kit for Illumina (NEB, cat.# E7770S). The NEBNext Adaptor for Illumina was diluted 1:5 (for 500 ng input RNA) or 1:25 (for 100 ng input RNA) in Tris/NaCl, pH 8.0 (10 mM Tris-HCl pH 8.0, 10 mM NaCl) and the ligation step was extended to 30 minutes. Libraries were enriched with 9-11 PCR cycles with the NEBNext Multiplex Oligos for Illumina (Dual Index Primers Set 1; NEB, cat.# E7600S). Library concentration was assessed by Qubit quantification (Qubit dsDNA HS Assay Kit; Invitrogen, cat.# Q32851). Multiplexed libraries were pooled and sequenced on the Illumina NovaSeq X Plus platform (150 bp, paired end reads) by MedGenome Inc. (Foster City, CA, USA).

##### RNAseq analysis

RNAseq raw reads were quality checked with FastQC (http://www.bioinformatics.babraham.ac.uk/projects/fastqc), trimmed with cutadapt (DOI: https://doi.org/10.14806/ej.17.1.200; version 4.5 with Python 3.10.13) and aligned onto the mouse genome (mm39) using STAR RNA-Seq aligner^107^ with the following options: --outSJfilterReads: Unique, --outFilterMultimapNmax: 1, -- outFilterIntronMotifs: RemoveNoncanonical, --outSAMstrandField: intronMotif. Samtools^108^ (version 1.9) was used to convert STAR output .sam files into .bam files, and to sort and index them. After counting how many reads overlapped an annotated gene (Ensembl GRCm39 annotations) using HTSeq^109^ (options: --htseq-count; -- stranded=no -f bam; --additional-attr=gene_name -m union), the output counts files were used to find DEGs with DESeq2^110^, run with default parameters within the Galaxy platform^111–113^. DEGs were called using an adjusted *p*-value ≤0.01, a fold change ≥2 and ≥10 mean counts. Gene transcript levels were visualized on the mm39 genome with the Integrative Genomics Viewer (IGV)^114,115^ using the bigWig output files from deepTools’ bamCoverage^116^ (version: 3.5.1; options: --binSize: 50; --extendReads: 250; --normalizeUsing: BPM; --samFlagInclude: 64). PCA and sample-to-sample distance analysis were part of the DESeq2 output.

Based on the DESeq2 output, pairwise comparison of conditions was performed by visualizing DEGs as called above via the EnhancedVolcano package (https://github.com/kevinblighe/EnhancedVolcano) within Bioconductor^117^ (release 3.20) using R (version 4.4.2) and indicating the number of up- or downregulated genes. In addition, DEGs of pairwise comparisons were further analyzed with respect to their enrichment in biological pathways by GO analysis using the clusterProfiler^118^ and AnnotationDbi (https://bioconductor.org/packages/AnnotationDbi) packages and selecting the ontology class BP for Biological processes.

Using the ComplexHeatmap package^119,120^, a comparison of DEGs in all five conditions was visualized in a heatmap displaying *z*-scores, whereby either all DEGs or selected DEGs were plotted. In addition, CPMs of some of these selected DEGs were plotted across all five conditions (barplot: mean of triplicates; CPM values of individual replicates as points).

#### Quantification and statistical analysis

The significance of the differences in the distribution of Sox9 expression levels (Fig. 3C), the distributions in cellular morphological features (Fig. 3D), as well as the percentage of proliferating Ki67-positive cells (Fig. 3E) and aberrant cell division phenotypes (Fig. S3B) was computed using a Mann-Whitney U test after aggregating all cells in one image and calculating a mean for each image. For Sox9 expression level analyses (Fig. 3B), alternative hypotheses in which cells in the Sox9 spheroid condition had stochastically greater metric values were used. For cellular morphologies (Fig. 3D), proliferation (Fig. 3E), and cell division phenotypes (Fig. S3B), a two-sided alternative hypothesis was used (Fig. 4D). *p*-values indicated in Fig. 3C-E and Fig. S3B were rated as follows: (ns) non-significant, *p*>0.05; (*) *p*≤0.05; (**) *p*≤0.01; (***) *p*≤0.001; (****) *p*≤0.0001. The exact *p*-values for pairwise comparisons in Fig. 3B-E are as follows: Fig. 3C left: *p*=0.05; Fig. 3C right: *p*=0.35; Fig. 3D left: *p*=7.2e- 5; Fig. 3D right: *p*=7.2e-5; Fig. 3E: *p*=0.01. The exact *p*-values for WT *vs.* Sox9_budding, WT *vs.* Sox9_spheroid, and Sox9_budding *vs.* Sox9_spheroid in Fig. S3B are as follows: Fig. S3B top left (a): 1.00, 1.95e-03, 3.07e-03; Fig. S3B top right (b): 5.55e-01, 4.91e-02, 2.06e-02; Fig. S3B bottom left (c): 7.84e-02, 1.01e-05, 6.70e-03; Fig. S3B bottom right (d): 7.57e-02, 1.11e-02, 2.35e-01.

For SMT experiments, bootstrapping on all combined experiments per fast SMT condition was performed by drawing 1000 samples from the population with replacement, whereby each sample contained as many nuclei as the total number of nuclei in the population under analysis. For each bootstrap replicate, diffusion spectra marginalized on the diffusion coefficients were generated and used to compute CIs for the fraction of bound trajectories. Code for SMT bootstrap analysis is included in our previously published^18^ custom-written Jupyter notebook (https://gitlab.com/tjian-darzacq-lab/walther_2dmsios_automatedsmt_2024/-/tree/main/SMT_analysis_scripts?ref_type=heads).

For determining a potential correlation between the relative POI expression level and the fraction bound in SMT experiments (Fig. 2D, Fig. 5D; Fig. S1G,H), nuclear Sox9 intensities were extracted from StarDist^102^ masks and plotted against the fraction bound of the POI in each nucleus inferred using saSPT’s^103^ State Array Dataset class for each POI. For each POI, a potential correlation between the fraction bound and the relative Sox9 expression level was calculated by performing a linear least-squares regression using the SciPy Python library^121^, which yielded an *R*-value (Pearson correlation coefficient). *p*-values were computed using a Wald test with a t-distribution test statistic and a two-sided alternative hypothesis. Code is included in our previously published^18^ custom-written Jupyter notebook (https://gitlab.com/tjian-darzacq-lab/walther_2dmsios_automatedsmt_2024/-/tree/main/SMT_analysis_scripts?ref_type=heads).

The corresponding figure legends contain statistical details for SMT experiments and fraction bound/intensity correlations mentioned above.

For PAPA-SMT experiments, statistical analyses were performed as described before using custom-written MATLAB scripts (^18,29^; https://github.com/tgwgraham/basic_PAPASMT_analysis). For a side-by-side comparison of the distributions for DR and PAPA trajectories, they were randomly subsampled without replacement for the condition with more trajectories. Following subsampling, bootstrapping analysis with replacement was performed on all combined experiments per condition for the PAPA datasets for Sox9_spheroid/Sox9, Sox9_spheroid/NLS, Sox9_budding/Sox9, and Sox9_budding/NLS. For each combined dataset, a random sample of size *n*, where *n* is the total number of cells in the combined dataset, was drawn 100 times. We reported the mean and standard deviation from these analyses (Fig. 6G,H; Fig. S7). For significance testing between DR and PAPA conditions, two-tailed *p*-values were calculated based on a normal distribution (SciPy^121^ function, scipy.stats.norm.sf) with mean equal to the difference between sample means and variance equal to the sum of the variances from the bootstrap resampling. The statistical details for the PAPA experiments shown in Fig. 6E-H are as follows: Sox9→Sox9: Subsampling of 9255 trajectories determined fractions bound to 40.3% (DR) and 52.0% (PAPA). Bootstrap resampling with 100 replicates determined fractions bound to 40.6±2.9% (DR) and 51.9±3.1% (PAPA); 2- sided *p*-value: 2.3e-7. Sox9→NLS: Subsampling of 4117 trajectories determined fractions bound to 26.4% (DR) and 26.1% (PAPA). Bootstrap resampling with 100 replicates determined fractions bound to 27.2±2.5% (DR) and 26.1±2.8% (PAPA); 2- sided *p*-value: 1.4. The statistical details for the PAPA experiments shown in Fig. S7 are as follows: Sox9→Sox9: Subsampling of 1191 trajectories determined fractions bound to 38.2% (DR) and 46.2% (PAPA). Bootstrap resampling with 100 replicates determined fractions bound to 38.9±4.9% (DR) and 46.0±6.6% (PAPA); 2-sided *p*- value: 0.09. Sox9→NLS: Subsampling of 2644 trajectories determined fractions bound to 16.8% (DR) and 16.3% (PAPA). Bootstrap resampling with 100 replicates determined fractions bound to 17.6±2.5% (DR) and 16.4±2.8% (PAPA); 2-sided *p*- value: 1.44. *p*-values indicated in Fig. 6G,H and Fig. S7 were rated as follows: (ns) non-significant *p*>0.05; (*) *p*≤0.05; (**) *p*≤0.01; (***) *p*≤0.001; (****) *p*≤0.0001.

---

**Figure S1:**
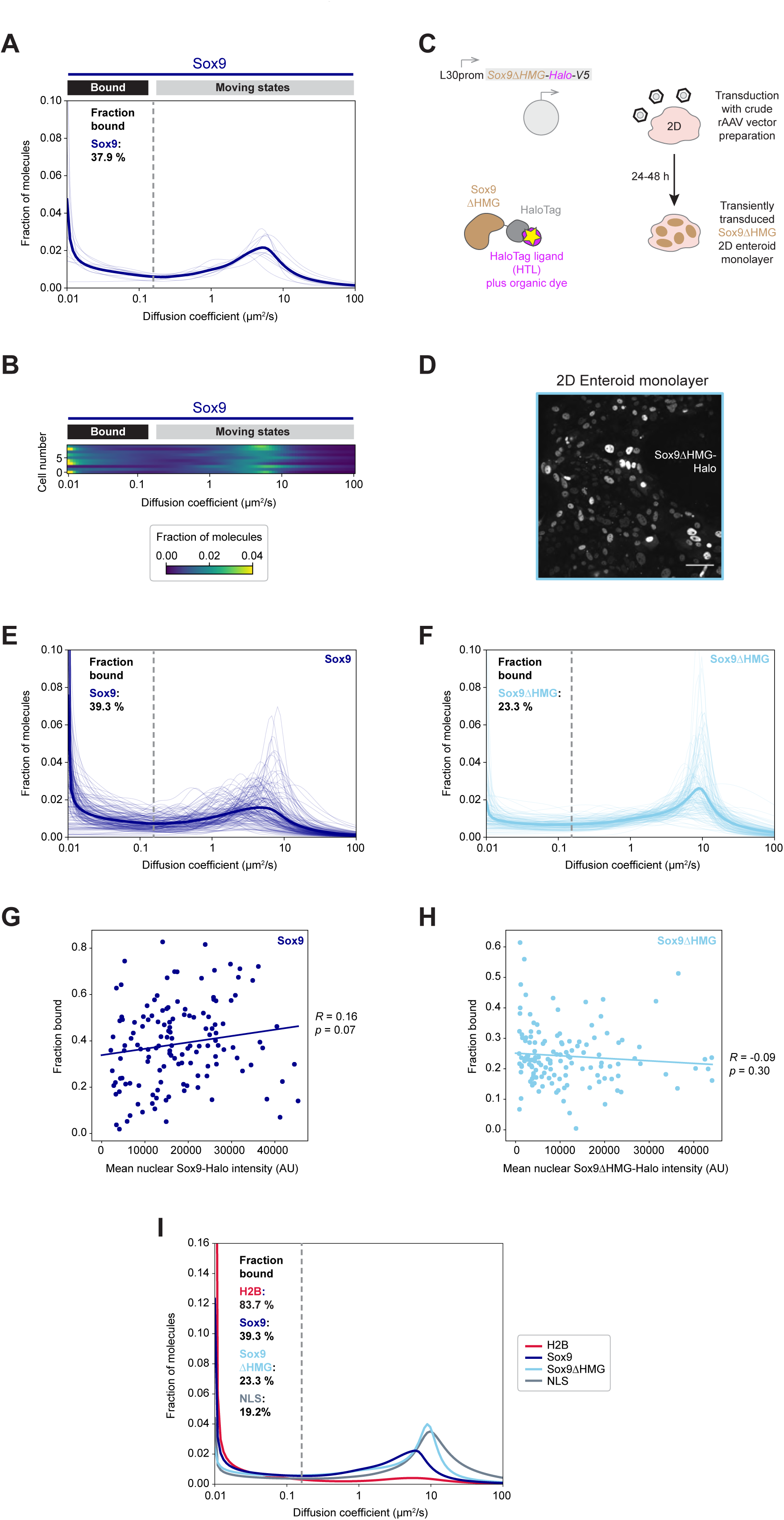
The diffusive behaviors of Sox9-Halo and its DNA binding domain mutant Sox9ΔHMG-Halo are expression level-independent, related to Figure 1. **(A,B)** Single-cell diffusion spectra (A) or diffusion heatmap (B) of a manual fast SMT experiment in Sox9-Halo 2D EMCs with a mean fraction bound of 37.9% (95% CI: 35.8-39.4%). In (B) cells are ordered by decreasing fraction bound from top to bottom (*n*=10 cells). **(C)** Generation of an organoid line transiently overexpressing a Sox9ΔHMG-HaloTag-V5 transgene through transduction with a crude rAAV vector preparation allows fluorescence detection (yellow star) of Sox9ΔHMG-Halo (brown- gray) covalently labeled with a dye-coupled HTL (magenta). **(D)** Confocal live imaging of a 2D EMC 48h post-transduction with a crude rAAV preparation to transiently express Sox9ΔHMG-Halo. Scale bar: 50 μm. **(E,F)** Single-cell diffusion spectra for 4 independent automated experiments for Sox9-Halo (E) and 5 independent manual experiments for Sox9ΔHMG-Halo (F). **(G,H)** Fractions bound for each cell are plotted against the mean nuclear fluorescence intensity for Sox9-Halo (G) or Sox9ΔHMG- Halo (H). Correlations between POI intensity and fraction bound (fitted line) were computed for each POI and the Pearson correlation coefficients (*R*) and *p*-values are indicated. **(I)** Mean diffusion spectra for H2B-Halo (red), Sox9-Halo (dark blue), Sox9ΔHMG-Halo (light blue), and Halo-NLS (gray). Bootstrap analysis of combined experiments with *n*=65,30,12,50,123,81,6 cells for H2B, *n*=10,44,42,56 cells for Sox9, *n*=16,42,34 cells for Sox9ΔHMG, and *n*=12,47,41 cells for NLS determined mean fractions bound of 83.7% (95% CI: 81.4-85.9%), 39.3% (95% CI: 32.9-46.7%), 23.3% (95% CI: 19.5-28.5%), and 19.2% (95% CI: 15.0-23.2%) respectively. Cells in (E-I) correspond to the 4 experiments for Sox9 (dark blue; combined *n*=152 cells) and the 5 experiments for Sox9ΔHMG (light blue; combined *n*=135 cells) plotted in Fig. 1E-I. Cells in (I) further correspond to the 7 experiments for H2B (red; combined *n*=367 cells) and the 3 experiments for NLS (gray; combined *n*=100 cells) plotted in Fig. 1G and Fig. S2H,I.

**Figure S2:**
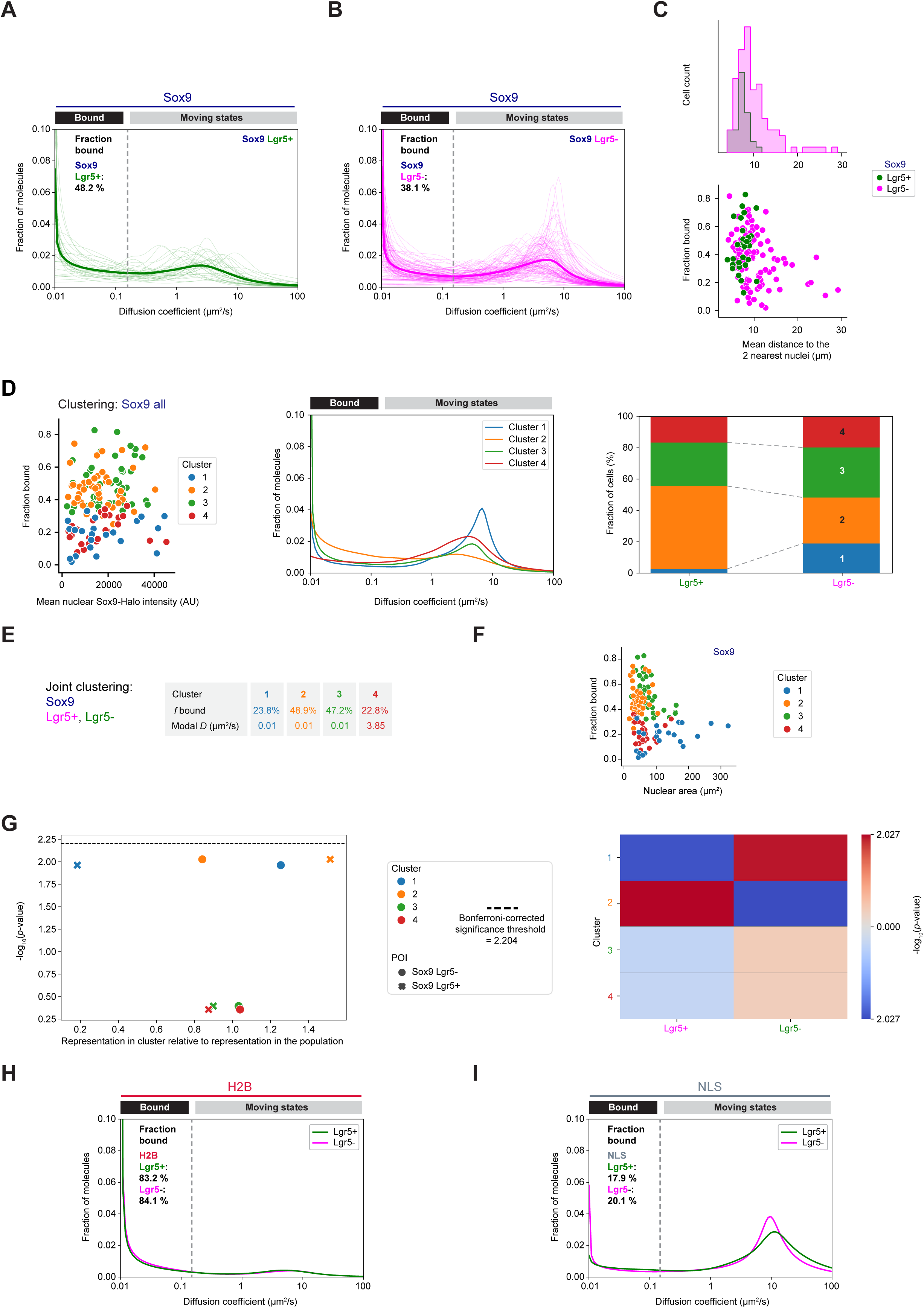
Differences in the diffusive behavior of Sox9-Halo in stem- and differentiated cell populations, but not for Halo-tagged immobile H2B and freely diffusing NLS CTRLs, related to Figure 2. **(A,B)** Single-cell diffusion spectra for 4 independent automated experiments for Sox9-Halo classified into Lgr5+ (green) (A) and Lgr5- (magenta) (B) subpopulations with *n*=36 and *n*=116 cells, respectively. **(C)** Top: Distribution of the single-cell morphological characteristic mean distance to the two nearest nuclei for Lgr5+/- (green/magenta) cells extracted from SMT data. Bottom: Single-cell correlation of SMT-derived fraction bound with mean distance to the two nearest nuclei for Lgr5+/- (green/magenta) cells. **(D-G)** Hierarchical clustering of all Sox9-Halo cells based on single-cell diffusion spectra using the Jensen-Shannon distance metric. (D) Left: Fractions bound for each cell plotted against the mean nuclear Sox9-Halo intensity with color-coded diffusion clusters. Middle: Mean diffusion spectra for each cluster. Right: Distribution of cells into diffusion clusters for Lgr5+/- (left/right). (E) Cluster statistics. (F) Fractions bound for each cell plotted against the nuclear area with color-coded diffusion clusters. (G) Left: *p*-values indicating the representation of Lgr5+ (x mark) and Lgr5- (circle) subpopulations in each diffusion cluster relative to the representation in the population; Bonferroni-corrected significance threshold (dashed line). Right: Heatmap of *p*-values indicating the representation of each subpopulation in each diffusion cluster (red: overrepresentation; blue: underrepresentation). Cluster: 1-blue, 2-orange, 3-green, 4- red. Shown are *n*=152 cells from 4 combined automated experiments (*n*=10,44,42,56 cells). **(H,I)** Mean diffusion spectra for Lgr5+/- (green/magenta) subpopulations of H2B-Halo (H) and Halo-NLS (I). Bootstrap analysis of combined experiments with *n*=65,30,12,50,123,81,6 cells for H2B and *n*=12,47,41 cells for NLS determined mean fractions bound of 83.2% (95% CI: 81.1-85.3%) and 84.1% (95% CI: 81.7-86.4%) as well as 17.9% (95% CI: 13.7-22.5%) and 20.1% (95% CI: 15.9-23.6%) for Lgr5+/- subpopulations, respectively. The Sox9 data are the same as in Fig.1E-I, Fig. S1E,G,I, Fig. 2B-D, and Fig. 5C. The H2B and NLS data are the same as in Fig. 1G and Fig. S1E.

**Figure S3:**
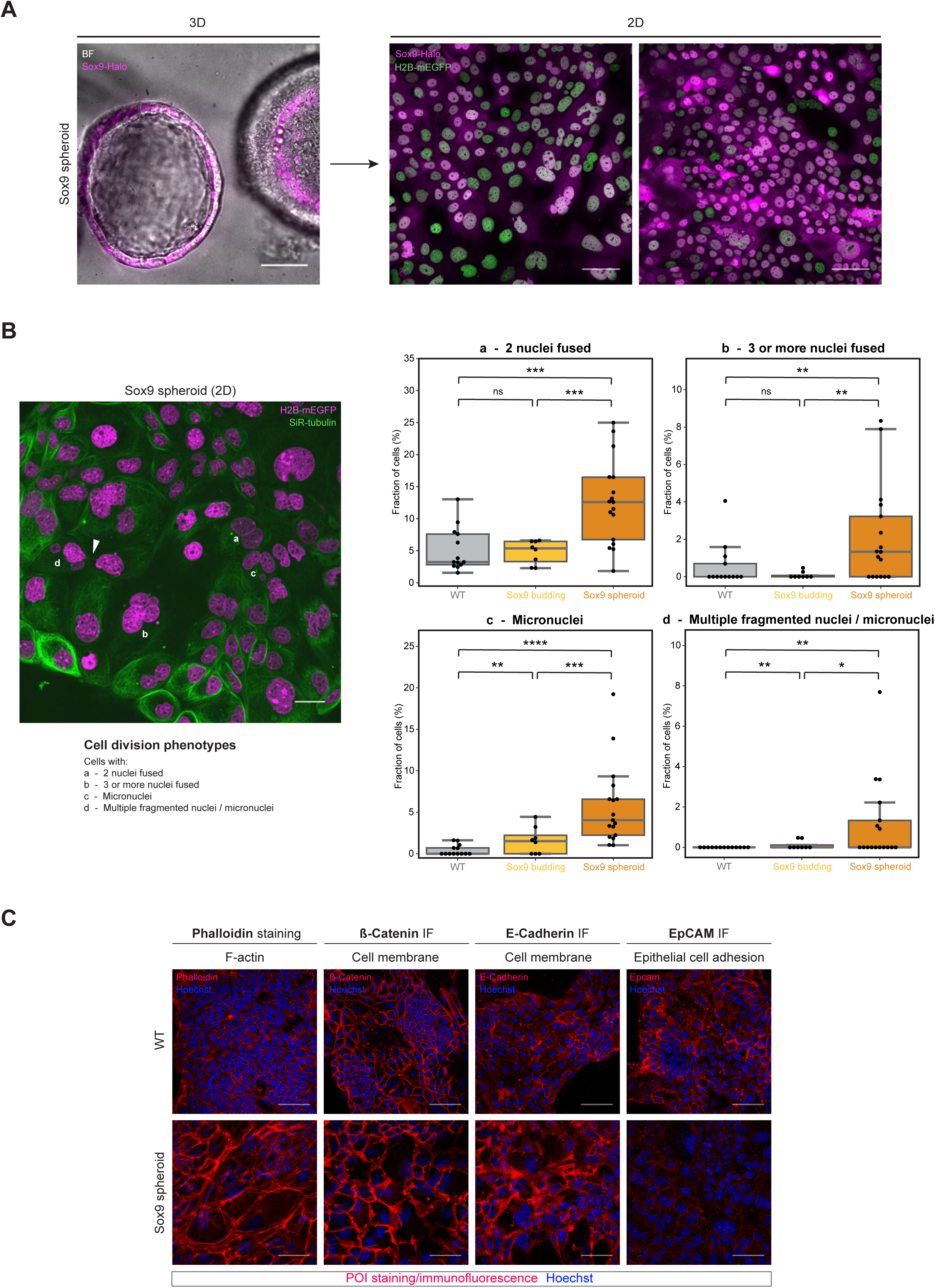
Sox9-Halo spheroids display cell division errors, actin stress fibers, and signs of cell membrane remodeling, related to Figure 3. **(A)** Confocal live imaging of Sox9-Halo spheroids co-expressing H2B-mEGFP 5 d post-seeding. Left: 3D spheroid (BF: gray; Sox9-Halo: magenta). Right: 2D EMC derived from 3D spheroid (Sox9-Halo: magenta; H2B-mEGFP: green). Scale bars: 50 μm. **(B)** Cell division phenotypes in 2D EMCs derived from Sox9-Halo spheroids. Left: Representative confocal image of live Sox9 spheroid-derived 2D EMC (H2B-mEGFP: magenta SiR-tubulin: green) with cell division phenotypes (a-d) and anaphase bridge (arrow) indicated. Scale bar: 20 μm. Right: Quantifications of cell division phenotypes (a: two nuclei fused; b: three or more nuclei fused; c: micronuclei; d: multiple fragmented nuclei or micronuclei) in 2D EMCs derived from WT (gray), Sox9- Halo_budding (yellow), or Sox9-Halo_spheroid (orange) organoids. Quantifications were based on 13 images with *n*=1821 cells for WT, 8 images with *n*=1626 cells for Sox9_budding, and 17 images with *n*=1756 cells for Sox9_spheroid. Each point represents one FOV; median: gray line, first/third quartile: whiskers; statistical testing based on Mann-Whitney U tests (see methods for details); (ns) non-significant, *p*>0.5; (*) *p*≤0.5; (**) *p*≤0.01; (***) *p*≤0.001; (****) *p*≤0.0001. **(C)** Confocal images of immunostained (POI: red) 2D EMCs derived from WT organoids (top) or Sox9-Halo spheroids (bottom) co-stained with Hoechst (blue). Scale bars: 50 μm.

**Figure S4:**
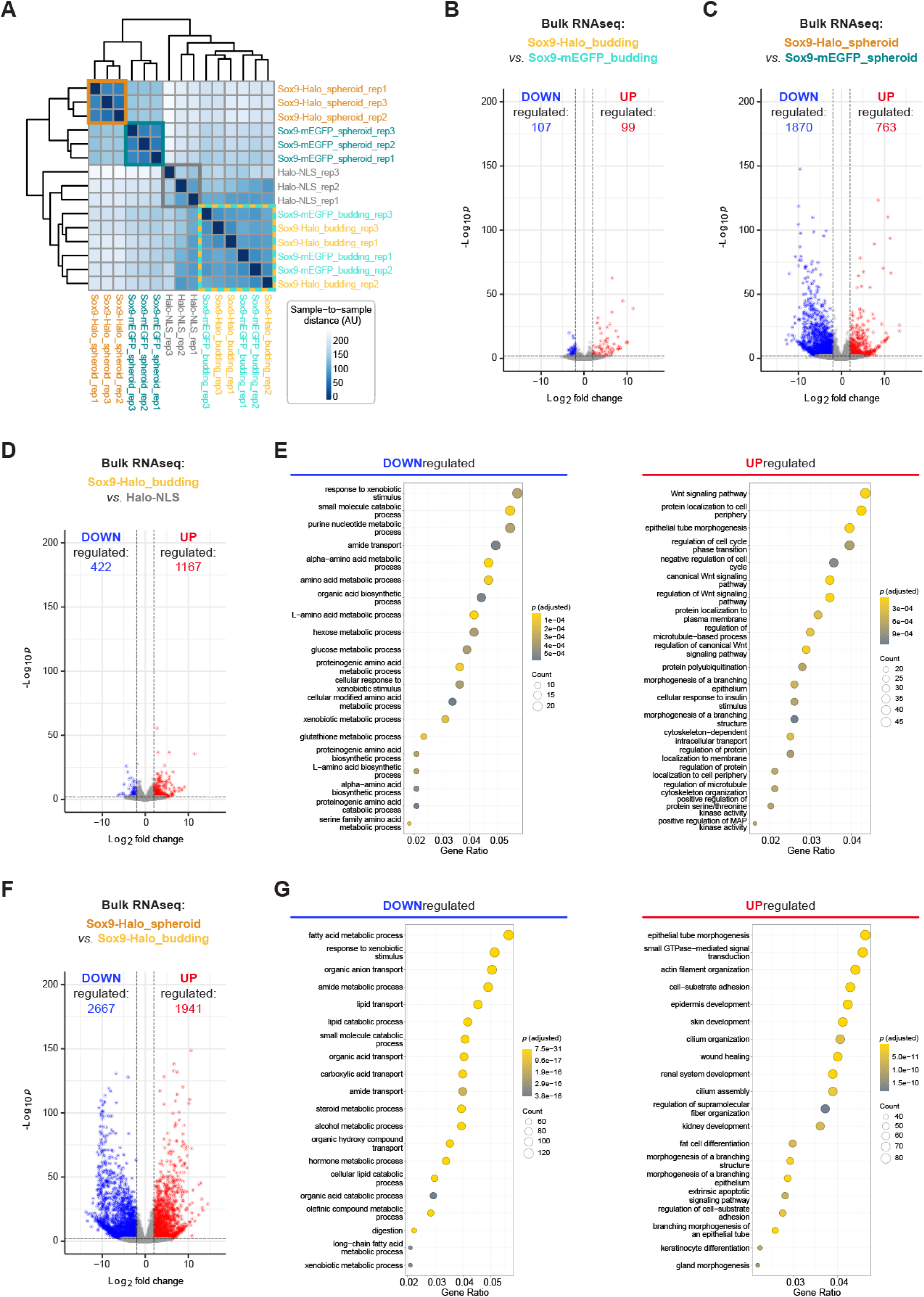
Enteroid reversion to a fetal-like state upon Sox9 overexpression is tag-independent and occurs through intermediate states with upregulated Wnt signaling-dependent pathways, related to Figure 4. **(A)** Sample-to-sample distances of the organoid samples Halo-NLS (gray), Sox9-mEGFP_budding (light turquoise), Sox9-Halo_budding (yellow), Sox9-mEGFP_spheroid (dark turquoise), and Sox9-Halo_spheroid (orange) in biological triplicates of bulk RNAseq experiments. **(B,C)** Volcano plot displaying DEGs (adjusted *p*-value ≤0.01, fold change ≥2 and mean counts ≥10; red/blue: up-/downregulated) in (B) Sox9-Halo_budding in comparison to Sox9-mEGFP_budding organoids or (C) Sox9-Halo spheroids in comparison to Sox9-mEGFP spheroids determined by bulk RNAseq. **(D)** Volcano plot displaying DEGs (adjusted *p*-value ≤0.01, fold change ≥2 and mean counts ≥10; red/blue: up-/downregulated) in Sox9-Halo_budding in comparison to Halo-NLS CTRL organoids determined by bulk RNAseq. **(E)** GO analysis for the top 20 biological pathways enriched in DEGs down- (left, blue) or upregulated (right, red) in Sox9- Halo_budding in comparison to Halo-NLS CTRL organoids with adjusted *p*-values and gene counts indicated. **(F)** Volcano plot displaying DEGs (adjusted *p*-value ≤0.01, fold change ≥2 and mean counts ≥10; red/blue: up-/downregulated) in Sox9-Halo spheroids in comparison to Sox9-Halo budding organoids determined by bulk RNAseq. **(E)** GO analysis for the top 20 biological pathways enriched in DEGs down- (left, blue) or upregulated (right, red) in Sox9-Halo spheroids in comparison to Sox9- Halo budding organoids with adjusted *p*-values and gene counts indicated. Data shown refers to the bulk RNAseq experiment from Fig. 4B-D and Fig. S5.

**Figure S5:**
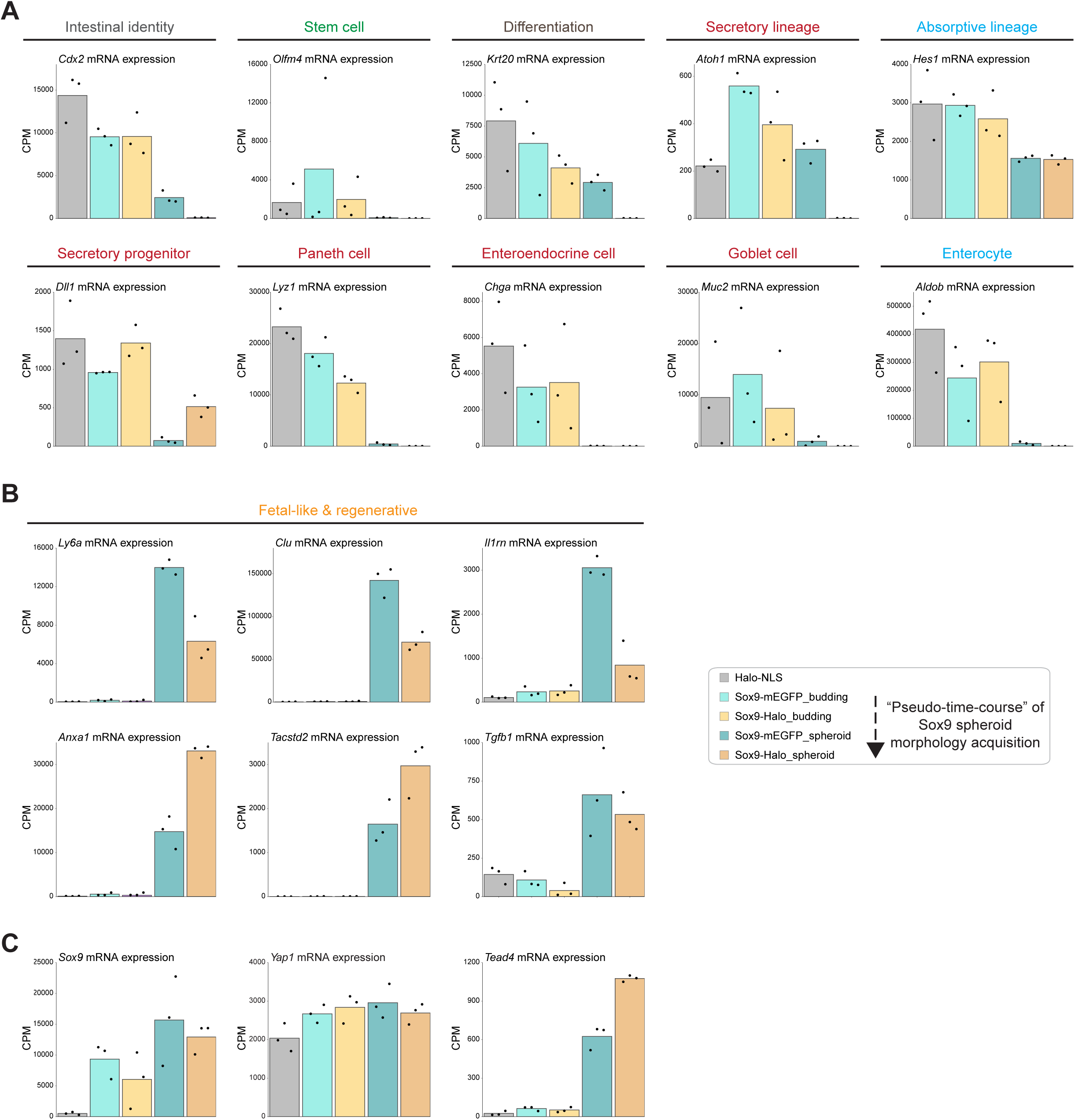
A “pseudo-time-course” of spheroid morphology acquisition upon Sox9 overexpression in enteroids reveals a reduction in intestinal epithelial signatures across lineages counteracted by the induction of a gene expression program resembling fetal-like reversion, which includes the upregulation of *Tead* acting downstream of Yap, related to Figure 4. (A-C) Selected DEGs in the stable organoid lines Halo-NLS (gray), Sox9-mEGFP_budding (light turquoise), Sox9- Halo_budding (yellow), Sox9-mEGFP_spheroid (dark turquoise), and Sox9- Halo_spheroid (orange) determined by bulk RNAseq in biological triplicates. The mean counts per million mapped reads (CPM) of three replicates (bar) and the CPMs for each replicate (points) are indicated. Samples are ordered according to passage number (time in culture) and degree of spheroid phenotype acquisition. (A) Intestinal identity, stem and differentiation markers of both secretory and absorptive lineages. (B) Regenerative fetal-like markers. (C) *Sox9*, *Yap1*, *Tead4*. Data shown refer to the bulk RNAseq experiment in Fig. 3F,G, Fig. 4B-D and Fig. S4.

**Figure S6:**
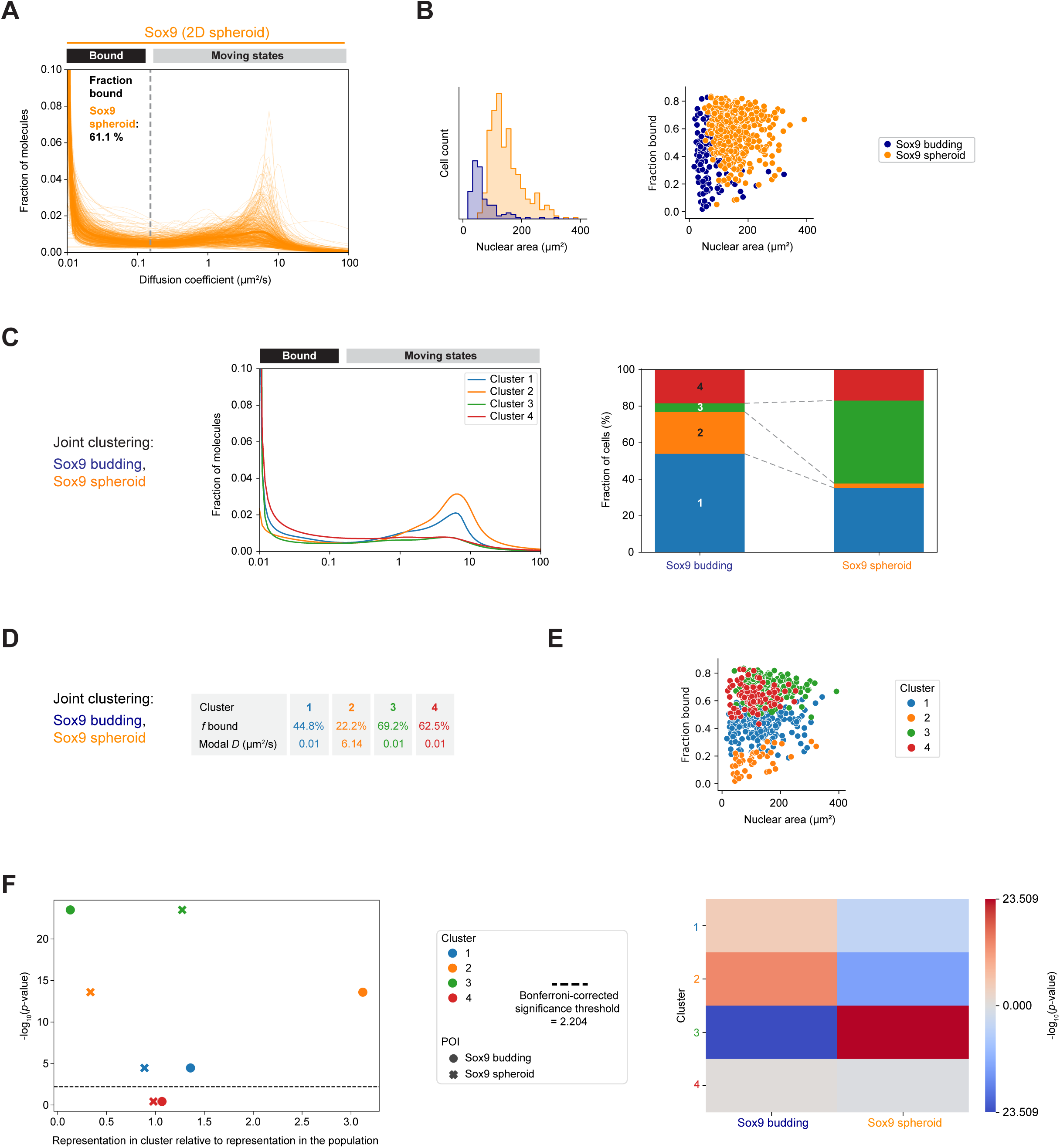
Unlike in budding enteroids, a larger fraction of immobile Sox9-Halo molecules is present in spheroid-derived EMCs with larger nuclei, related to Figure 5. **(A)** Single-cell diffusion spectra for 10 independent automated experiments for Sox9-Halo in spheroid-derived 2D EMCs. **(B)** Left: Nuclear area distribution for Sox9-Halo_budding (dark blue) and Sox9-Halo_spheroid (orange) cells determined from SMT data in 2D EMCs. Right: Single-cell correlation of SMT-derived fraction bound with nuclear area for Sox9-Halo_budding (dark blue) and Sox9-Halo_spheroid (orange). **(C-F)** Joint hierarchical clustering of Sox9-Halo_budding and Sox9- Halo_spheroid based on single-cell diffusion spectra using the Jensen-Shannon distance metric. (C) Left: Mean diffusion spectra of each cluster. Right: Distribution of cells from both Sox9-Halo samples into diffusion clusters. (D) Cluster statistics. (E) Fractions bound for each cell against the nuclear area with color-coded diffusion clusters. (F) Left: *p*-values indicating the representation of Sox9-Halo_budding (circle) and Sox9-Halo_spheroid (x mark) in each diffusion cluster relative to the representation in the population; Bonferroni-corrected significance threshold (dashed line). Right: Heatmap of *p*-values indicating the representation of each sample in each diffusion cluster (red: overrepresentation; blue: underrepresentation). Cluster: 1-blue, 2-orange, 3-green, 4-red. The Sox9-Halo_budding data are the same as in Fig. 1E-I, Fig. 2B-D, Fig. 5C, Fig. S1E,G,I, and Fig. S2A-G. The Sox9-Halo_spheroid data are the same as in Fig. 5B-D.

**Figure S7:**
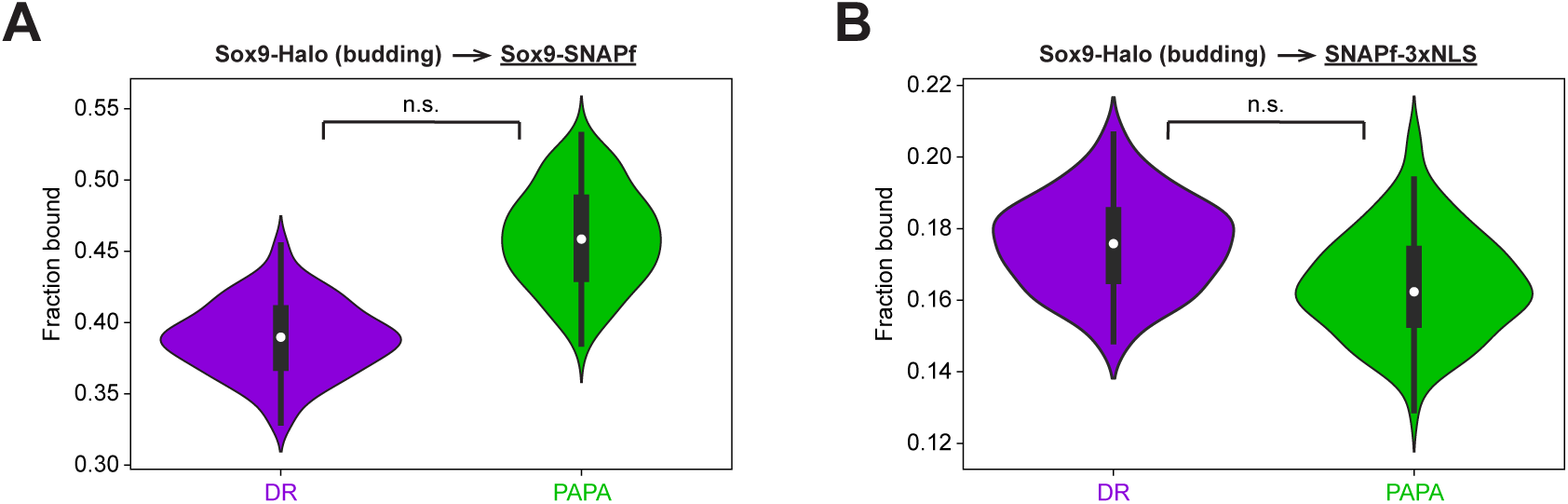
PAPA-SMT reveals a chromatin-bound pool of self-associated Sox9 in live 2D EMCs derived from budding Sox9-Halo enteroids, related to Figure 6. **(A,B)** PAPA experiments in 2D EMCs derived from Sox9-Halo_budding organoids. Violin plots for fractions bound (median: white point; first/third quartile: whiskers) of Sox9-Halo→Sox9-SNAPf (A) and Sox9-Halo→SNAPf-3xNLS (B) determined from DR/PAPA (purple/green) trajectories. Data in (A) are from 6 combined experiments with *n*=108 cells (*n*=6,25,32,7,29,9 cells; 1661 DR and 1191 PAPA trajectories; bootstrapped fractions bound: 38.9±4.9% (DR) and 46.0±6.6% (PAPA)). Data in (B) are from 5 combined experiments with *n*=97 cells (*n*=22,44,2,14,15 cells; 3579 DR and 2644 PAPA trajectories; bootstrapped fractions bound: 17.6±2.5% (DR) and 16.4±2.8% (PAPA)). (n.s.) *p*>0.05. For statistical details see methods.

